# Predicting Alzheimer’s disease progression using deep recurrent neural networks

**DOI:** 10.1101/755058

**Authors:** Minh Nguyen, Tong He, Lijun An, Daniel C. Alexander, Jiashi Feng, B.T. Thomas Yeo, for the Alzheimer’s Disease Neuroimaging Initiative

## Abstract

Early identification of individuals at risk of developing Alzheimer’s disease (AD) dementia is important for developing disease-modifying therapies. In this study, given multimodal AD markers and clinical diagnosis of an individual from one or more timepoints, we seek to predict the clinical diagnosis, cognition and ventricular volume of the individual for every month (indefinitely) into the future. We proposed and applied a minimal recurrent neural network (minimalRNN) model to data from The Alzheimer’s Disease Prediction Of Longitudinal Evolution (TADPOLE) challenge, comprising longitudinal data of 1677 participants (Marinescu et al. 2018) from the Alzheimer’s Disease Neuroimaging Initiative (ADNI). We compared the performance of the minimalRNN model and four baseline algorithms up to 6 years into the future. Most previous work on predicting AD progression ignore the issue of missing data, which is a prevalent issue in longitudinal data. Here, we explored three different strategies to handle missing data. Two of the strategies treated the missing data as a “preprocessing” issue, by imputing the missing data using the previous timepoint (“forward filling”) or linear interpolation (“linear filling). The third strategy utilized the minimalRNN model itself to fill in the missing data both during training and testing (“model filling”). Our analyses suggest that the minimalRNN with “model filling” compared favorably with baseline algorithms, including support vector machine/regression, linear state space (LSS) model, and long short-term memory (LSTM) model. Importantly, although the training procedure utilized longitudinal data, we found that the trained minimalRNN model exhibited similar performance, when using only 1 input timepoint or 4 input timepoints, suggesting that our approach might work well with just cross-sectional data. An earlier version of our approach was ranked 5th (out of 53 entries) in the TADPOLE challenge in 2019. The current approach is ranked 2nd out of 63 entries as of June 3rd, 2020.

## 1 Introduction

Alzheimer’s disease (AD) dementia is a devastating neurodegenerative disease with a long prodromal phase and no available cure. It is widely believed that an effective treatment strategy should target individuals at risk for AD early in the disease process (Scheltens et al., 2016). Consequently, there is significant interest in predicting the longitudinal disease progression of individuals. A major difficulty is that although AD commonly presents as an amnestic syndrome, there is significant heterogeneity across individuals (Murray et al., 2011; Noh et al., 2014; Zhang et al., 2016; Risacher et al., 2017; Young et al., 2018; Sun et al., 2019). Since AD dementia is marked by beta-amyloid- and tau-mediated injuries, followed by brain atrophy and cognitive decline (Jack et al., 2010, 2013), a multimodal approach might be more effective than a single modality approach to disentangle this heterogeneity and predict longitudinal disease progression (Marinescu et al., 2018, 2020).

In this study, we proposed a machine learning algorithm to predict multimodal AD markers (e.g., ventricular volume, cognitive scores, etc) and clinical diagnosis of individual participants for every month up to six years into the future. Most previous work has focused on a “static” variant of the problem, where the goal is to predict a single timepoint (Duchesne et al., 2009; Stonnington et al., 2010; Zhang and Shen, 2012; Moradi et al., 2015; Albert et al., 2018; Ding et al., 2018) or a set of *pre-specified* timepoints in the future (regularized regression; (Wang et al., 2012; Johnson et al., 2012; McArdle et al., 2016; Wang et al., 2016)). By contrast, our goal is the longitudinal prediction of clinical diagnosis and multimodal AD markers at a potentially unlimited number of timepoints into the future^1^, as defined by The Alzheimer’s Disease Prediction Of Longitudinal Evolution (TADPOLE) challenge (Marinescu et al., 2018, 2020), which arguably a more relevant and complete goal for tasks, such as prognosis and cohort selection.

One popular approach to this longitudinal prediction problem is mixed-effect regression modeling, where longitudinal trajectories of AD biomarkers are parameterized by linear or sigmoidal curves (Vemuri et al., 2009; Ito et al., 2010; Sabuncu et al., 2014; Samtani et al., 2012; Zhu and Sabuncu, 2018). However, such a modeling approach requires knowing the shapes of the biomarker trajectories a priori. Furthermore, even though the biomarker trajectories might be linear or sigmoidal when *averaged* across participants (Caroli and Frisoni, 2010; Jack et al., 2010; Sabuncu et al., 2011), individual subjects might deviate significantly from the assumed parametric forms.

Consequently, it might be advantageous to not assume that the biomarker trajectories follow a specific functional form. For example, Xie and colleagues proposed an incremental regression modeling approach to predict the next timepoint based on a fixed number of input time points (Xie et al., 2016). The prediction can then be used as input to predict the next timepoint and so on indefinitely. However, the training procedure requires participants to have two timepoints, thus “wasting” data from participants with less or more than two timepoints. Therefore, state-based models that do not constrain the shapes of the biomarker trajectories or assume a fixed number of timepoints might be more suitable for this longitudinal prediction problem (e.g., discrete state hidden Markov models; Sukkar et al. 2012). Here, we considered recurrent neural networks (RNNs), which allow an individual’s latent state to be represented by a vector of numbers, thus providing a richer encoding of an individual’s “disease state” beyond a single integer (as in the case of discrete state hidden Markov models). In the context of medical applications, RNNs have been used to model electronic health records (Lipton et al., 2016a; Choi et al., 2016; Esteban et al., 2016; Pham et al., 2017; Rajkomar et al., 2018; Suo et al., 2018) and AD disease progression (Nguyen et al., 2018; Ghazi et al., 2019).

Most previous work on predicting AD progression ignore the issue of missing data (Stonnington et al., 2010; Sukkar et al., 2012; Lei et al., 2017; Liu et al., 2019). However, missing data is prevalent in real-world applications and arises due to study design, delay in data collection, subject attrition or mistakes in data collection. Missing data poses a major difficulty for modeling longitudinal data since most statistical models assume feature-complete data (García-Laencina et al., 2010). Many studies sidestep this issue by removing subjects or timepoints with missing data, thus potentially losing a large quantity of data. There are two main approaches for handling missing data (Schafer and Graham 2002). First, the “preprocessing” approach handles the missing data issue in a separate preprocessing step, by imputing the missing data (e.g., using the missing variable’s mean or more sophisticated machine learning strategies; Azur et al., 2011; Rehfeld et al., 2011; Stekhoven and Bühlmann, 2011; White et al., 2011; Zhou et al., 2013), and then using the imputed data for subsequent modeling. Second, the “integrative” approach is to integrate the missing data issue directly into the models or training strategies, e.g., marginalizing the missing data in Bayesian approaches (Marquand et al., 2014; Wang et al., 2014; Aksman et al., 2019).

In this work, we proposed to adapt the minimalRNN model (Chen, 2017) to predict AD progression. The minimalRNN has fewer parameters than other RNN models, such as the long short-term memory (LSTM) model, so it might be less prone to overfitting. Although RNNs are usually trained using feature-complete data, we explored two “preprocessing” and one “integrative” approaches to deal with missing data. We used data from the TADPOLE competition, comprising longitudinal data from 1677 participants (Marinescu et al. 2018; 2019). An earlier version of this work was published at the International Workshop on Pattern Recognition in Neuroimaging and utilized the more complex LSTM model (Nguyen et al., 2018). Here, we extended our previous work by using a simpler RNN model, expanding our comparisons with baseline approaches and exploring how the number of input timepoints affected prediction performance. We also compared the original LSTM and current minimalRNN models using the live leaderboard on TADPOLE.

## 2 Methods

### 2.1 Problem setup

The problem setup follows that of the TADPOLE challenge (Marinescu et al. 2018). Given the multimodal AD markers and diagnostic status of a participant from one or more timepoints, we seek to predict the cognition (as measured by ADAS-Cog13; Mohs et al., 1997), ventricular volume (as measured by structural MRI) and clinical diagnosis of the participant for every month indefinitely into the future.

### 2.2 Data

We utilized the data provided by the TADPOLE challenge (Marinescu et al., 2018). The data consisted of 1677 subjects from the ADNI database (Jack et al., 2008). Each participant was scanned at multiple timepoints. The average number of timepoints was 7.3 ± 4.0 (Figure 1A), while the average number of years from the first timepoint to the last timepoint was 3.6 ± 2.5 (Figure 1B).

**Figure 1.**
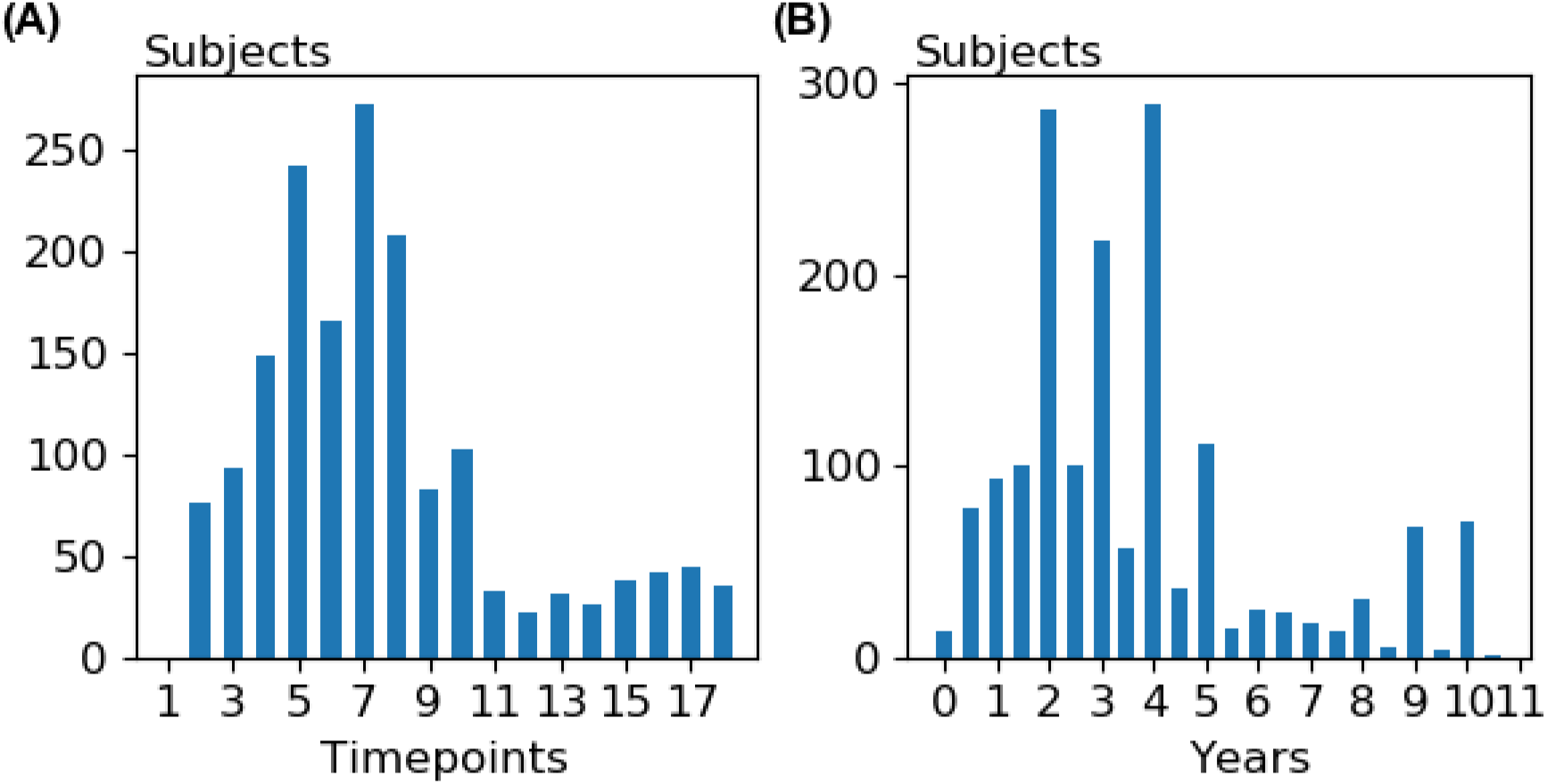
**(A)** Distribution of the number of timepoints for all subjects in the dataset. **(B)** Distribution of the number of years between the first and last timepoints for all subjects in the dataset.

For consistency, we used the same set of 23 variables recommended by the TADPOLE challenge, which included diagnosis, neuropsychological test scores, anatomical features derived from T1 magnetic resonance imaging (MRI), positron emission tomography (PET) measures and CSF markers (Table 1). The diagnostic categories corresponded to normal control (NC), mild cognitive impairment (MCI) and Alzheimer’s disease (AD).

**Table 1.** Set of variables together with their means, standard deviations and percentage of timepoints where the variables were actually observed. SB: Sum of boxes, ADAS: Alzheimer’s Disease Assessment Scale, RAVLT: Rey Auditory Verbal Learning Test

| | mean (± std) | % timepoints with measures |
| --- | --- | --- |
| Clinical Dementia Rating Scale (SB) | $2.17 \pm 2.81 \times 10^0$ | 70.36 % |
| ADAS-Cog11 | $1.13 \pm 0.86 \times 10^1$ | 69.95 % |
| ADAS-Cog13 | $1.75 \pm 1.16 \times 10^1$ | 69.27 % |
| Mini-Mental State Examination (MMSE) | $2.65 \pm 0.39 \times 10^1$ | 70.12 % |
| RAVLT immediate | $3.44 \pm 1.36 \times 10^1$ | 69.33 % |
| RAVLT learning | $4.02 \pm 2.81 \times 10^0$ | 69.33 % |
| RAVLT forgetting | $4.23 \pm 2.52 \times 10^0$ | 69.12 % |
| RAVLT forgetting percent | $5.97 \pm 3.83 \times 10^1$ | 68.57 % |
| Functional Activities Questionnaire (FAQ) | $5.59 \pm 7.92 \times 10^0$ | 70.60 % |
| Montreal Cognitive Assessment (MOCA) | $2.30 \pm 0.47 \times 10^1$ | 38.99 % |
| Ventricles | $4.21 \pm 2.32 \times 10^4$ | 58.44 % |
| Hippocampus | $6.68 \pm 1.24 \times 10^3$ | 53.39 % |
| Whole brain volume | $1.01 \pm 0.11 \times 10^6$ | 60.35 % |
| Entorhinal cortical volume | $3.44 \pm 0.81 \times 10^3$ | 50.78 % |
| Fusiform cortical volume | $1.71 \pm 0.28 \times 10^4$ | 50.78 % |
| Middle temporal cortical volume | $1.92 \pm 0.31 \times 10^4$ | 50.78 % |
| Intracranial volume | $1.53 \pm 0.16 \times 10^6$ | 62.43 % |
| Florbetapir (18F-AV-45) - PET | $1.19 \pm 0.22 \times 10^0$ | 16.62 % |
| Fluorodeoxyglucose (FDG) - PET | $1.20 \pm 0.16 \times 10^0$ | 26.31 % |
| Beta-amyloid (CSF) | $1.02 \pm 0.59 \times 10^3$ | 18.60 % |
| Total tau | $2.93 \pm 1.30 \times 10^2$ | 18.55 % |
| Phosphorylated tau | $4.80 \pm 1.44 \times 10^1$ | 18.62 % |
| Diagnosis | - | 69.89 % |

### 2.3 Proposed model

We adapted the minimalRNN (Chen, 2017) for predicting disease progression. Here, we utilized minimalRNN instead of LSTM because it has less parameters and is therefore less likely to overfit (see Appendix A for details). The model architecture and update equations are shown in Figure 2. Let ***x***_***t***_ denote all variables observed at time *t*, comprising the diagnosis ***s***_***t***_ and remaining continuous variables ***g***_***t***_ (Eq. 1 in Figure 2B). Here, diagnosis was represented using one-hot encoding. In other words, diagnosis was represented as a vector of length three. More specifically, if the first entry was one, then the participant was a normal control. If the second entry was one, then the participant was mild cognitively impaired. If the third entry was one, then the participant had AD dementia. For now, we assume that all variables were observed at all timepoints; the missing data issue will be addressed in Section 2.4.

**Figure 2.**
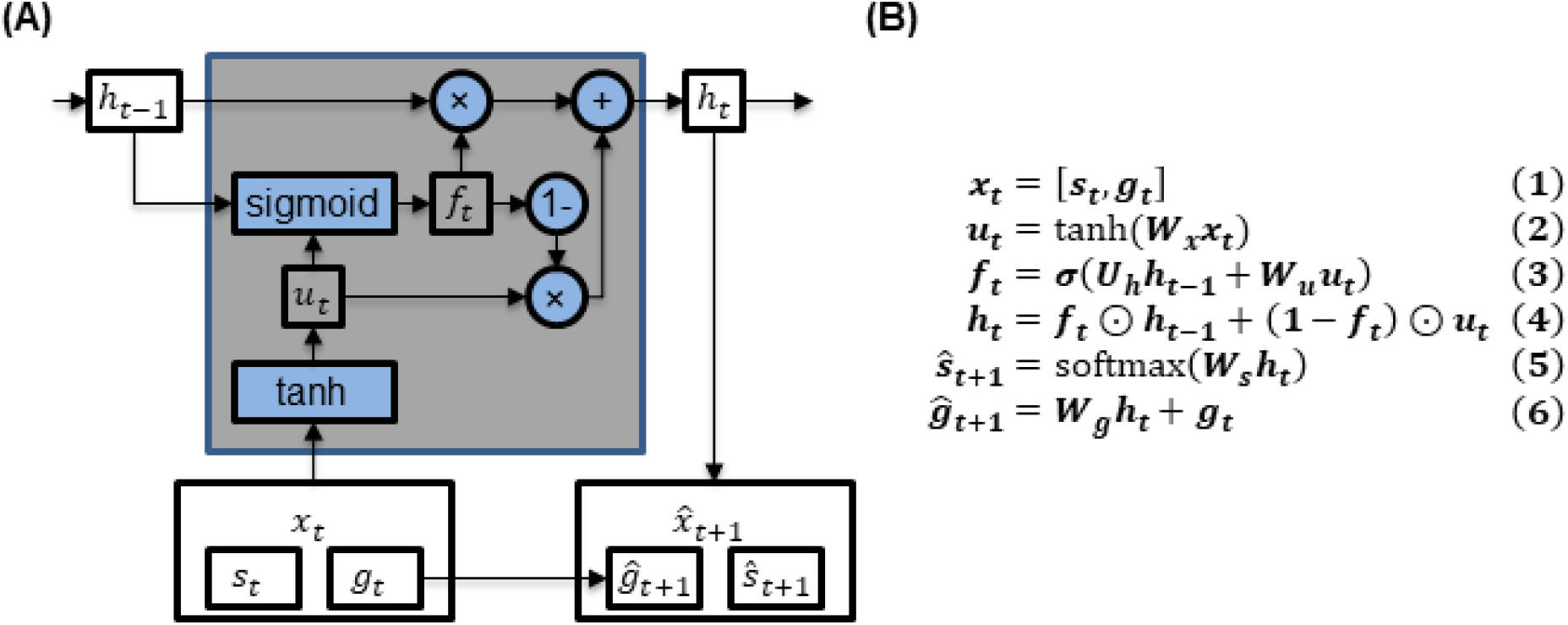
**(A)** MinimalRNN. **(B)** MinimalRNN update equations. ***s***_***t***_ and ***g***_***t***_ denote categorical (i.e., diagnosis) and continuous variables respectively (Table 1). The input ***x***_***t***_ to each RNN cell comprised the diagnosis ***s***_***t***_ and continuous variables ***g***_***t***_ (Eq. 1). Note that ***s***_***t***_ was represented using one-hot encoding. The hidden state ***h***_***t***_ was a combination of the previous hidden state ***h***_***t***−**1**_ and the transformed input ***u***_***t***_ (Eq. 4). The forget gate ***f*** _***t***_ weighed the contributions of the previous hidden state ***h***_***t***−**1**_ and current transformed input ***u***_***t***_ toward the current hidden state ***h***_***t***_ (Eq. 3). The model predicted the next month diagnosis 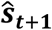 and continuous variables 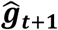 using the hidden state ***h***_***t***_ (Eqs. 5 and 6). ⊙ and **σ** denote element-wise product and the sigmoid function respectively.

At each timepoint, the transformed input ***u***_***t***_ (Eq. 2 in Figure 2) and the previous hidden state ***h***_***t***−**1**_ were used to update the hidden state ***h***_***t***_ (Eqs. 3 and 4 in Figure 2B). The hidden state can be interpreted as integrating all information about the subject up until that timepoint. The hidden state ***h***_***t***_ was then used to predict the observations at the next timepoint ***x***_***t***+**1**_ (Eqs. 5 and 6 in Figure 2B).

In the ADNI database, data were collected at a minimum interval of 6 months. However, in practice, data might be collected at an unscheduled time (e.g., month 8 instead of month 6). Consequently, the duration between timepoints *t* and *t* + 1 in the RNN was set to be 1 month. However, experiments with different durations were also performed with little impact on the results (see Section 2.7.2).

#### 2.3.1 Training with no missing data

The RNN training is illustrated in Figure 3. The RNN was trained to predict the next observation (***x***_***t***_) given the previous observations (***x***_**1**_, ***x***_**2**_, …, ***x***_***t***−**1**_). The errors between the predicted outputs (e.g. 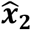) and the ground truth outputs (e.g. ***x***_**2**_) were used to update the model parameters. The error (or loss *L*) was defined as follows:

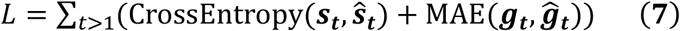

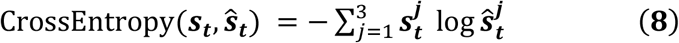

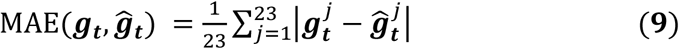

**Figure 3.**
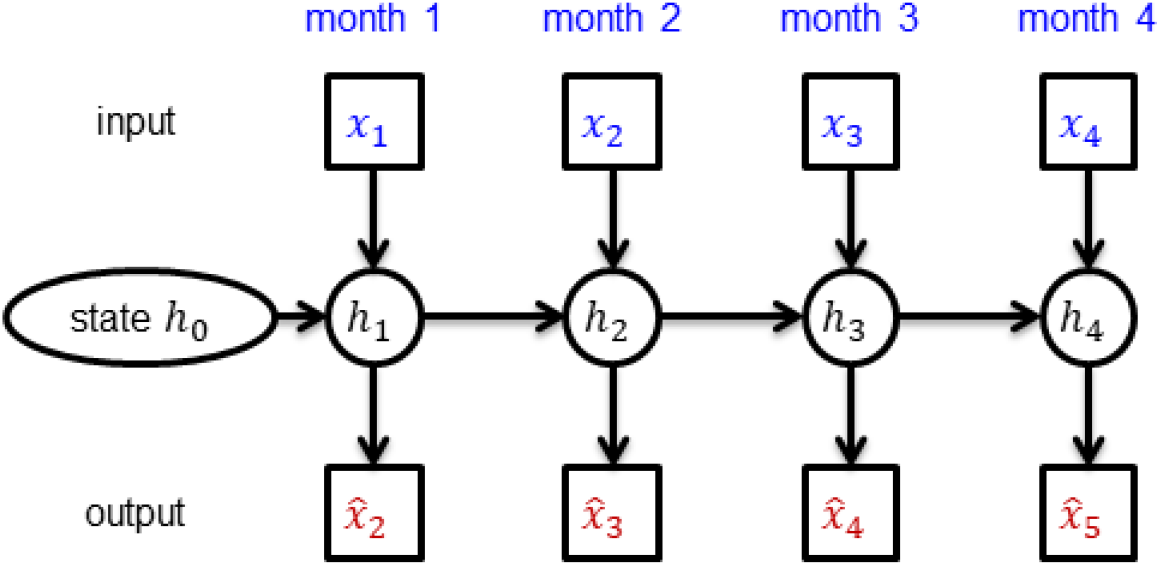
The minimalRNN was trained to predict the next observation given the current observation (e.g., predicting 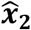 given ***x***_**1**_). Errors between the actual observations (e.g., ***x***_**2**_) and predictions (e.g., 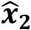) were used to update the model parameters. The hidden state ***h***_***t***_ encoded information about the subject up until time *t*.

It is important to note that the loss function was only evaluated using available observations. Missing data were not considered when computing the loss. Furthermore, we note that the two terms in the loss function (Eq. 7) were weighted equally. Changing the relative weights of the two terms could potentially influence the model performance. However, this would increase the number of hyperparameters, so we did not experiment with varying the weighting in this study. The value of ***h***_**0**_ was set to be **0**. During training, gradients of loss *L* with respect to the model parameters were back-propagated to update the RNN parameters. The RNN was trained using Adam (Kingma and Ba, 2015).

#### 2.3.2 Prediction with no missing data

Figure 4 illustrates how the RNN was used to predict AD progression in an example subject (from the validation or test set). Given observations for months 1, 2 and 3, the goal of the model was to predict observations in future months. From month 4 onwards, the model predictions (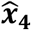 and 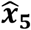) were fed in as inputs to the RNN (for months 5 and 6 respectively) to make further predictions (dashed lines in Figure 4).

**Figure 4.**
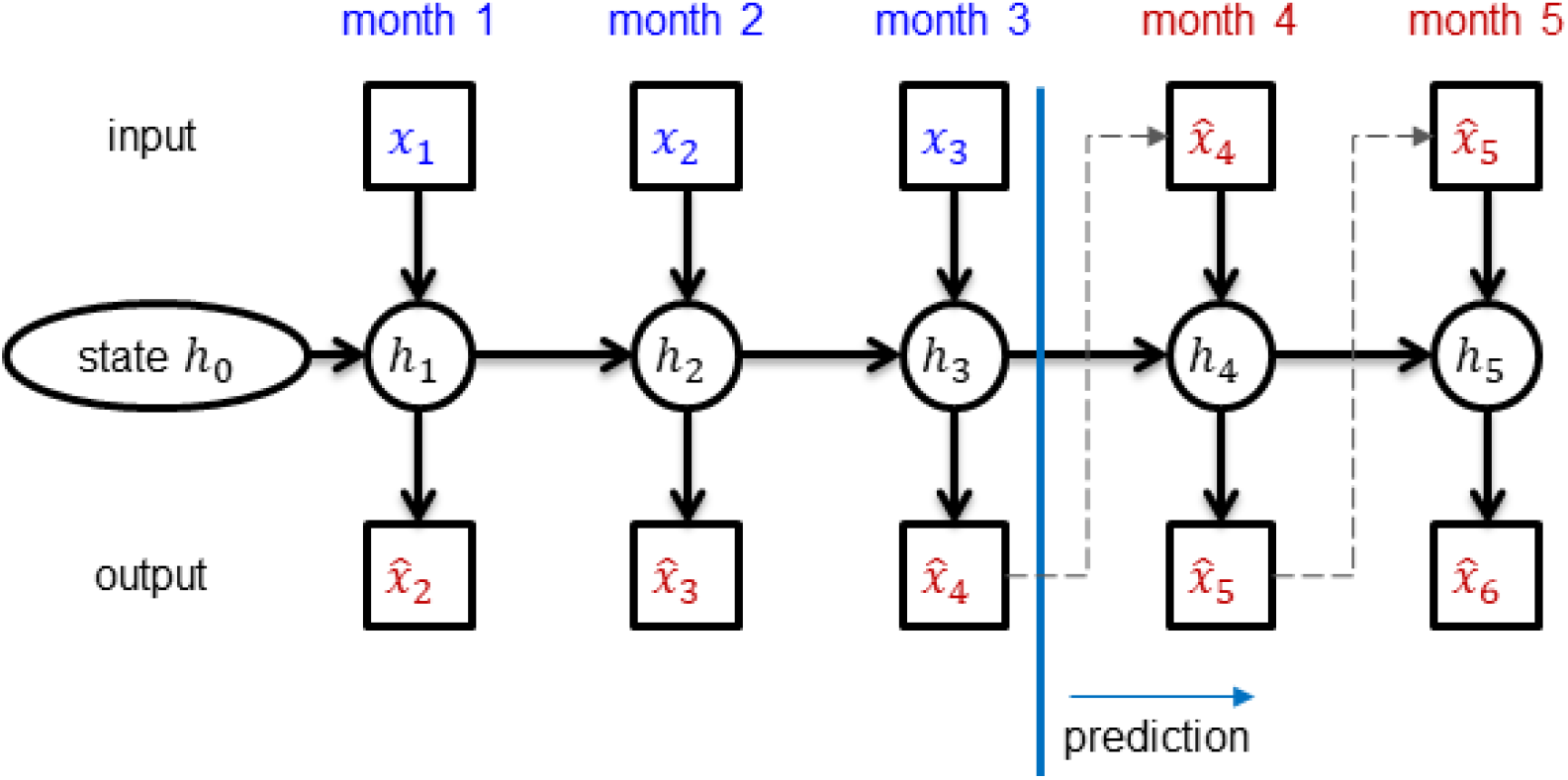
Predicting future timepoints (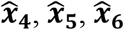, etc) given three initial timepoints (***x***_**1**_, ***x***_**2**_, and ***x***_**3**_). Prediction started at month 4. Since there were no observed data at timepoints 4 and 5, the predictions (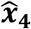 and 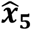) were used as inputs (at timepoints 5 and 6 respectively) to predict further into the future.

### 2.4 Missing data

As seen in Table 1, there were a lot of missing data in ADNI. This was exacerbated by the fact that data were collected at a minimum interval of 6 months, while the sampling period in the RNN was set to be 1 month (to handle off-schedule data collection). During training, the loss function was evaluated only at timepoints with available observations. Similarly, when evaluating model performance (Section 2.6), only available observations were utilized.

The missing data also posed a problem for the RNN update equations (Figure 2B), which assumed all variables were observed. Here, we explored two “preprocessing” strategies (Sections 2.4.1 & 2.4.2) and one “integrative” strategy (Section 2.4.3) to handle the missing values. As explained in the introduction, “preprocessing” strategies impute the missing data in a separate preprocessing. The imputed data is then used for subsequent modeling. On the other hand, “integrative” strategies incorporate the missing data issue directly into the model or training strategies.

#### 2.4.1 Forward filling

Forward filling involved imputing the data using the last timepoint with available data (Che et al., 2018; Lipton et al., 2016b). Figure 5A illustrates an example of how forward-filling in time was used to fill in missing input data. In this example, there were two input variables A and B. The values of feature A at time t = 2, 3 and 4 were filled using the last observed value of feature A (at time t = 1). Similarly, the values at t = 7, 8 of feature A were filled using value at t = 6 when it was last observed. If data was missing at the first timepoint, the mean value across all timepoints of all training subjects was used for the imputation.

**Figure 5.**
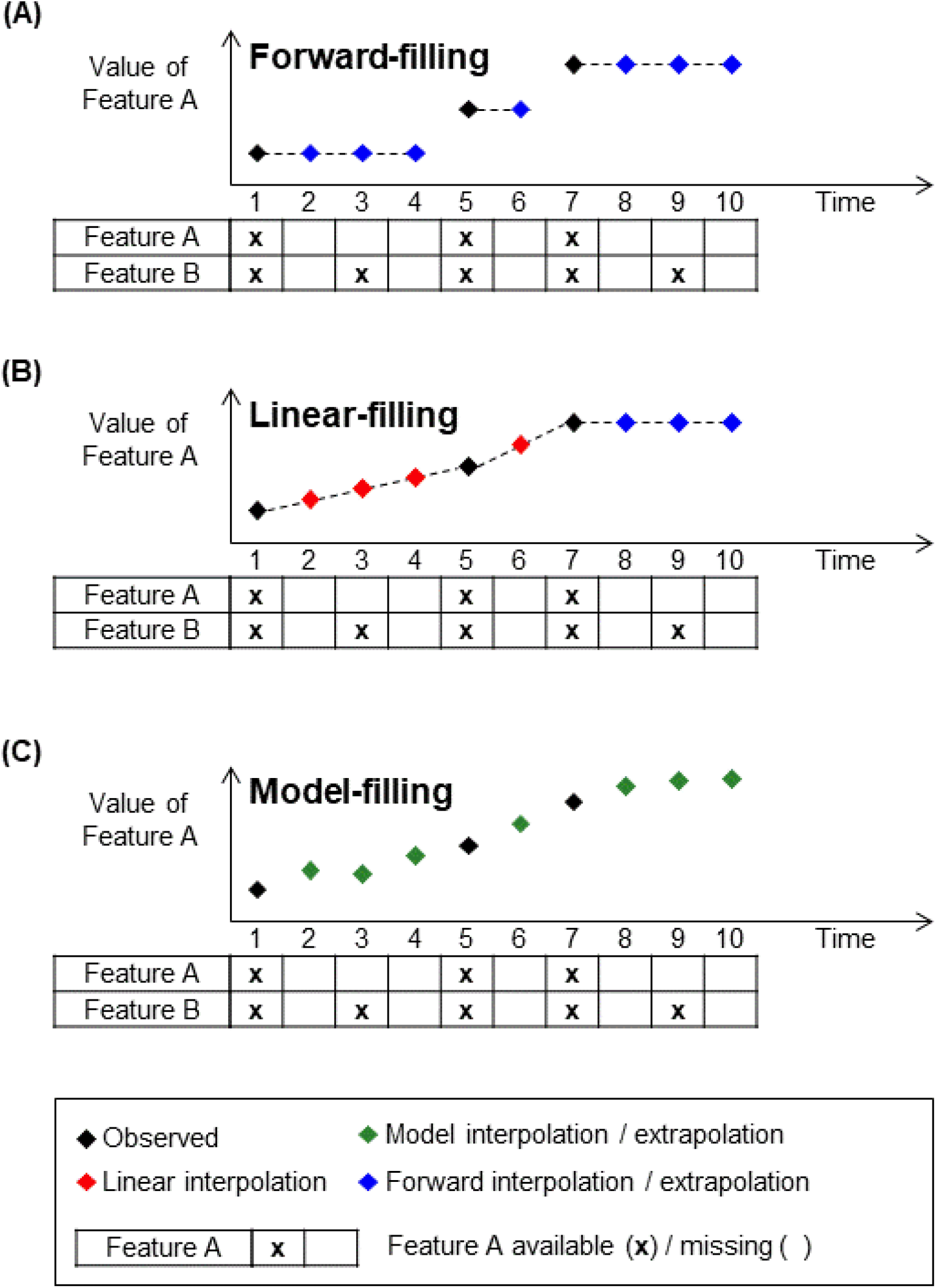
Different strategies to impute missing data. (A) Forward-filling imputed missing values using the last observed value. (B) Linear-filling imputed missing values using linear interpolation between previous observed and next observed values. Notice that linear-filling did not work for months 8, 9 and 10 because there was no future observed data for linear interpolation, so forward filling was utilized for those timepoints. (C) Model-filling imputed missing values using model predictions.

#### 2.4.2 Linear filling

The previous strategy utilized information from previous timepoints for imputation. One could imagine that it might be helpful to use previous and future timepoints for imputation. The linear filling strategy performed linear interpolation between the previous timepoint and the next time point with available data (Junninen et al., 2004). Figure 5B shows an example of linear interpolation. Values of feature A at time t = 2, 3, 4, 6 were filled in using linear interpolation. However, linear-filling did not work for months 8, 9 and 10 because there was no future observed data for linear interpolation, so forward-filling was utilized for those timepoints. Like forward filling, if data was missing at the first timepoint, the mean value across all timepoints of all training subjects was used for the imputation.

#### 2.4.3 Model filling

We also considered a novel model filling strategy of filling in missing data. As seen in Section 2.3.2 (Figure 5), the prediction of the RNN could be used as inputs for the next timepoint. The same approach can be used for filling in missing data.

Figure 5C shows an example of how the RNN was used to fill in missing data. At time t = 2 to 6, the values of feature A were filled in using predictions from the RNN. The RNN could also be used to extrapolate features that “terminate early” (e.g., time t = 8 and 9).

A theoretical benefit of modeling filling was that the full sets of features were utilized for the imputation. For example, both features A and B at time t = 1 were used by the RNN to predict both input features at time t = 2 (Figure 5C). This was in contrast to forward or linear filling, which would utilize only feature A (or B) to impute feature A (or B).

Like forward filling, if data was missing at the first timepoint, the mean value across all timepoints of all training subjects was used for the imputation.

### 2.5 Baselines

We considered four baselines: constant prediction, support vector machine/regression (SVM/SVR), linear state-space (LSS) model, and long short-term memory (LSTM) model.

#### 2.5.1 Constant prediction

The constant prediction algorithm simply predicted all future values to be the same as the last observed values. The algorithm did not need any training. While this might seem like an overly simplistic algorithm, we will see that the constant prediction algorithm is quite competitive for near term prediction.

#### 2.5.2 SVM/SVR

As explained in the introduction, most previous studies have focused on a “static” variant of the problem, where the goal is to predict a single timepoint or a set of pre-specified timepoints in the future. Here, we will consider such a baseline by using SVM to predict clinical diagnosis (which was categorical) and SVR to predict ADAS-Cog13 and ventricular volume (which were continuous). The models were implemented using scikit-learn (Pedregosa et al., 2011). We note that separate models were trained for each target variable (clinical diagnosis, ADAS-Cog13 and ventricular volume).

Because SVM/SVR accepts fixed length feature vectors, it cannot handle subjects with different number of input timepoints. Therefore, we trained different SVM/SVR models using 1 to 4 input timepoints (spaced 6 months apart) to predict the future. The 6-month interval was chosen because the ADNI data was collected roughly every 6 months. As can be seen in Section 3.1, the best results were obtained with 2 or 3 input timepoints, so we did not explore more than 4 input timepoints. The features were concatenated across the input timepoints. For example, since there were 23 features at each timepoint, then for the “2 input timepoints” SVM/SVR models, the input features constituted a vector of length 46. On the other hand, for the “3 input timepoints” SVM/SVR models, the input features constituted a vector of length 69.

For each SVM/SVR baseline, we trained separate SVM/SVR models to predict 10 sets of timepoints (spaced 6 months apart) into the future, i.e., 6, 12, 18, …, 60 months into the future. 60 months were the maximum because of insufficient data to train SVM/SVR to predict further into the future (Figure 1B). To summarize, separate SVM/SVR models were trained for different target variable (clinical diagnosis, ADAS-Cog13 and ventricular volume), for different number of input timepoints (1, 2, 3 or 4 input timepoints) and for different number of future predictions (6, 12, 18, …, 60 months). This yielded a total of 3 × 4 × 10 = 120 SVM/SVR models.

To maximize the number of data samples for training, we used all available timepoints in the training subjects to train the SVM/SVR models. For example, let us consider a training subject with 10 observed timepoints spaced 6 months apart. In the case of the SVM/SVR models with one input timepoint, this subject would contribute 9 training samples to train a model for predicting 6 months ahead, 8 training samples to train a model for predicting 12 months ahead, 7 training samples to train a model for predicting 18 months ahead, and so on.

The linear filling strategy (Figure 5B) was used to handle missing data. We also experimented with using multivariate functional principal component analysis (MFPCA) for filling in the missing data (Happ and Greven, 2018; Li et al., 2018). Because prediction performance was evaluated at every month in the future, prediction at intermediate months (e.g., months 1 to 5, months 7 to 11, etc) were linearly interpolated. Prediction from month 61 onwards utilized forward filling based on the prediction at month 60.

One tricky issue arose when a test subject had insufficient input timepoints for a particular SVM/SVR baseline. For example, the 4-timepoint SVM/SVR baseline required 4 input timepoints in order to predict future timepoints. In this scenario, if a test subject only had 2 input timepoints, then the 2-timepoint SVM/SVR was utilized for this subject even though we were considering the 4-timepoint SVM/SVR baseline. We utilized this strategy (instead of discarding the test subject) in order to ensure the test sets were exactly the same across all algorithms.

#### 2.5.3 Linear state space (LSS) model

We considered a linear state space (LSS) baseline by linearizing the minimalRNN model (Figure 6). Other than the update equations (Figure 6), all other aspects of training and prediction were kept the same. For example, the LSS models utilized the same data imputation strategies (Section 2.4) and were trained with the same cost function using Adam.

**Figure 6.**
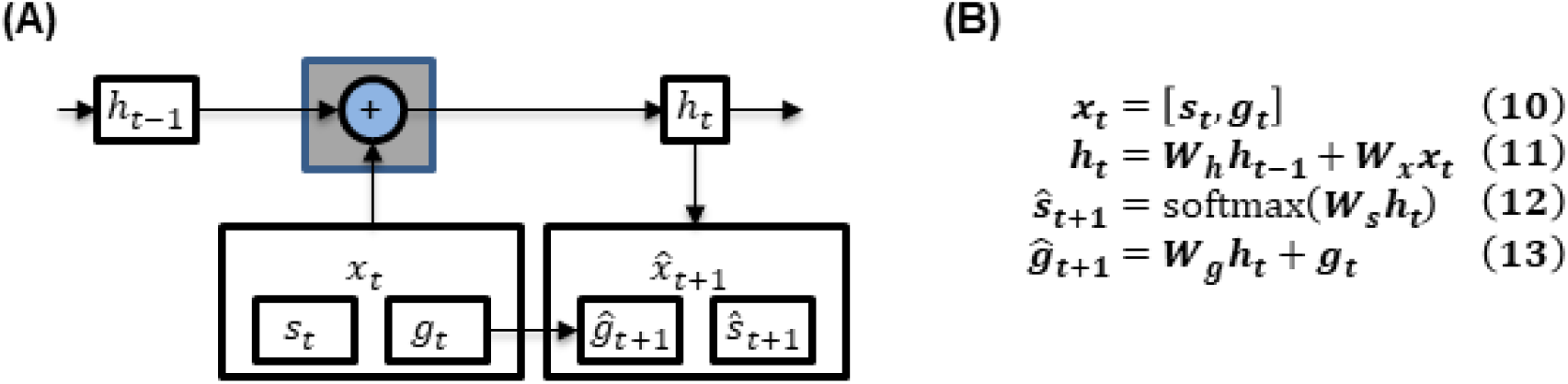
**(A)** Linear state space (LSS) model. Observe the gray cell is much simpler than the minimalRNN **(B)** LSS update equations. ***s***_***t***_ and ***g***_***t***_ denote categorical (i.e., diagnosis) and continuous variables respectively (Table 1). The input ***x***_***t***_ to each LSS cell comprised the diagnosis ***s***_***t***_ and continuous variables ***g***_***t***_ (Eq. 10). Like before, ***s***_***t***_ was represented using one-hot encoding. The hidden state ***h***_***t***_ was a combination of the previous hidden state ***h***_***t***−**1**_ and the input ***x***_***t***_ (Eq. 11). The model predicted the next month diagnosis 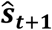 and continuous variables 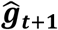 using the hidden state ***h***_***t***_ (Eqs. 12 and 13).

#### 2.5.4 Long Short Term Memory (LSTM) model

The LSTM model is widely used for modelling sequences and temporal trajectories (Ghazi et al., 2019; Lipton et al., 2016a). We have previously used LSTM for predicting AD progression (Nguyen et al., 2018). Here, we favor minimalRNN over LSTM models, as they have less parameters, so are less prone to overfitting when data is limited. See Appendix A for further discussion.

### 2.6 Performance evaluation

We randomly divided the data into training, validation and test sets. The ratio of subjects in the training, validation and test sets was 18:1:1. The training set was used to train the model. The validation set was used to select the hyperparameters. The test set was used to evaluate the models’ performance. For subjects in the validation and test sets, the first half of the timepoints of each subject were used to predict the second half of the timepoints of the same subject. All variables (except diagnostic category, which was categorical rather than continuous) were z-normalized. The z-normalization was performed on the training set. The mean and standard deviation from the training set was then utilized to z-normalize the validation and test sets. The random split of the data into training, validation and test sets was repeated 20 times to ensure stability of results (Kong et al., 2019; Li et al., 2019; Varoquaux, 2018). Care was taken so that the test sets were non-overlapping so that the test sets across the 20 data splits covered the entire dataset.

The HORD algorithm (Regis and Shoemaker 2013; Eriksson, Bindel, and Shoemaker 2015; Ilievski et al. 2017) was utilized to find the best hyperparameters by maximizing model performance on the validation set. We note that this optimization was performed independently for each training/validation/test split of the dataset. The hyperparameter search space for minimalRNN, LSS, and LSTM is shown in Table 2. The hyperparameter search space for the SVM/SVR is shown in Table 3. The final set of hyperparameters are found in Tables S1 to S13.

**Table 2.** Hyperparameter search space of MinimalRNN, LSS and LSTM estimated from the validation sets using HORD.

| Hyper-parameter | Range |
| --- | --- |
| Input dropout rate | 0.0 – 0.5 |
| Recurrent dropout rate | 0.0 – 0.5 |
| L2 weight regularization | $10^{-7} - 10^{-5}$ |
| Learning rate | $10^{-5} - 10^{-2}$ |
| Number of hidden layers | 1 – 3 |
| Size of hidden state | 128 – 512 |

**Table 3.** Hyperparameter search space of the SVM/SVR models estimated from the validation sets using HORD.

|  | SVM | SVR |
| --- | --- | --- |
| Kernel | Linear or RBF |  |
| Epsilon | NA | $10^{-3} - 10^{-0}$ |
| Penalty | $10^{-3} - 10^3$ | |
| Gamma | $10^{-3} - 10^3$ | |

Following the TADPOLE competition, diagnosis classification accuracy was evaluated using the multiclass area under the operating curve (mAUC; Hand and Till, 2001) and balanced class accuracy (BCA) metrics. The mAUC was computed as the average of three two-class AUC (AD vs not AD, MCI vs not MCI, and CN vs not CN). For both mAUC and BCA metrics, higher values indicate better performance. ADAS-Cog13 and ventricles prediction accuracy was evaluated using mean absolute error (MAE). Lower MAE indicates better performance. The final performance for each model was computed by averaging the results across the 20 test sets. Even though the 20 test sets do not overlap, the subjects used for training the models do overlap across the test sets. Therefore, the prediction performances were not independent across the 20 test sets. To account for the non-independence, we utilized the corrected resampled t-test (Bouckaert and Frank, 2004) to evaluate differences in performance between models.

### 2.7 Further analysis

#### 2.7.1 Impact of the number of input timepoints on prediction accuracy

For the minimalRNN to be useful in clinical settings, it should ideally be able to perform well with as little input timepoints as possible. Therefore, we applied the best model (Section 2.6) to the test subjects using only 1, 2, 3 or 4 input timepoints (Figure 7). This is different from the main benchmarking analysis (Section 2.6), where all input timepoints (which accounted for half of the total number of timepoints) of the test subjects were used for predicting future timepoints. Test subjects with less than 4 input timepoints were discarded, so that the same test subjects were evaluated across the four conditions (i.e., 1, 2, 3 or 4 input timepoints). Because we discarded some test subjects, the result of this analysis is not comparable to that of the main benchmarking analysis (Section 2.6).

**Figure 7.**
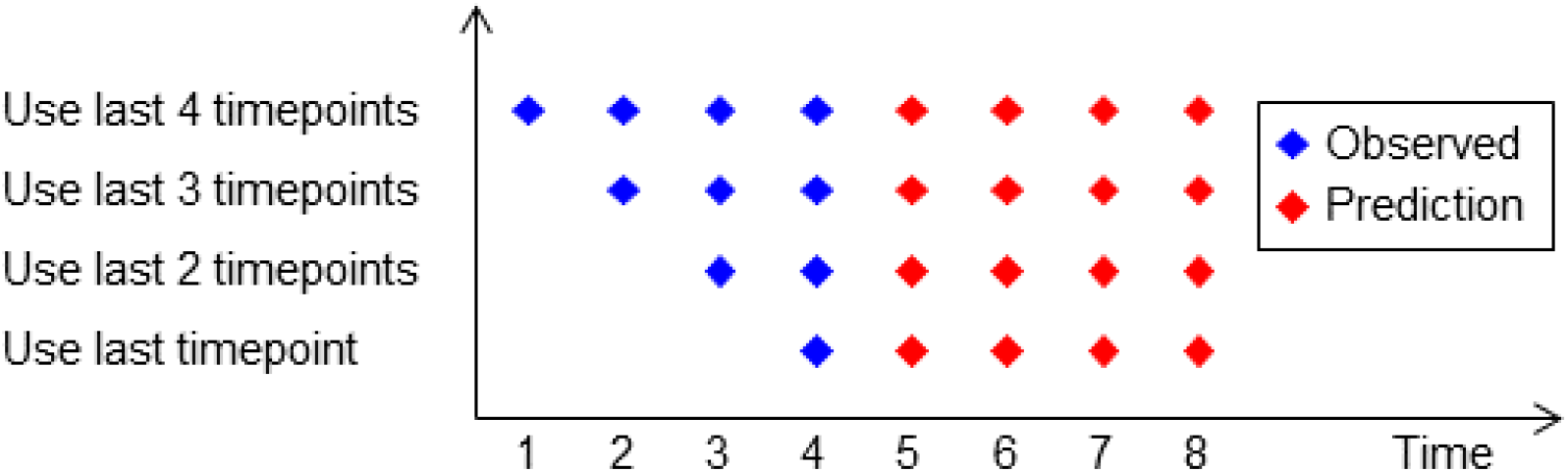
Prediction performance as a function of the number of input timepoints in the test subjects.

#### 2.7.2 Effect of temporal resolution of minimalRNN

Even though the ADNI data was collected at a minimum interval of 6 months, in practice, data was not collected at exactly 6-month interval, e.g., the data might be collected at month 4, instead of the scheduled data collection at month 6. Furthermore, the TADPOLE challenge required participants to make future prediction at a monthly interval with prediction performance evaluated at a monthly resolution. Therefore, our main analysis utilized minimalRNN models with a temporal resolution of 1 month.

However, the choice of temporal resolution (i.e., number of months between timepoints) might affect the performance of the minimalRNN. For example, using a finer temporal resolution (e.g., 1-month interval versus 6-month interval) leads to more missing data, which might lead to worse performance. On the other hand, using a coarser temporal resolution (e.g., 6-month interval versus 1-month interval) leads to greater mis-alignment between the minimalRNN’s timepoints and the actual observations. For example, if we consider a minimalRNN with a temporal resolution of 6 months, then actual observed data at month 10 would need to be assigned to month 12, which might lead to worse performance. Finally, using a coarser temporal resolution results in fewer hidden state updates between two points in time, making it potentially easier for the minimalRNN to learn longer-term temporal patterns.

Here, we experimented with three different temporal resolutions: 1-month interval, 3-month interval, and 6-month interval. The RNN models were trained and tested using the same procedure described in Section 2.6, including hyperparameter search. For training the 3-month and 6-month minimalRNN models, observed data were assigned to the closest timepoint. To evaluate performance of the 3-month and 6-month minimalRNN models, their predictions were linearly interpolated to obtain a temporal resolution of 1 month. Performance was evaluated only at timepoints with observed ground truth data.

#### 2.7.3 Impact of different terms in the minimalRNN model

To investigate which term in the minimalRNN model is important for model performance, we conducted ablation experiments whereby we gradually simplify the MinimalRNN update equations in 4 steps (Figure 8). In the last step (Variant 4), the simplified update equations were the same as the update equations of the linear state space (LSS) model. The ablated RNN models were trained and tested using the same procedure described in Section 2.6, including hyperparameter search.

**Figure 8.**
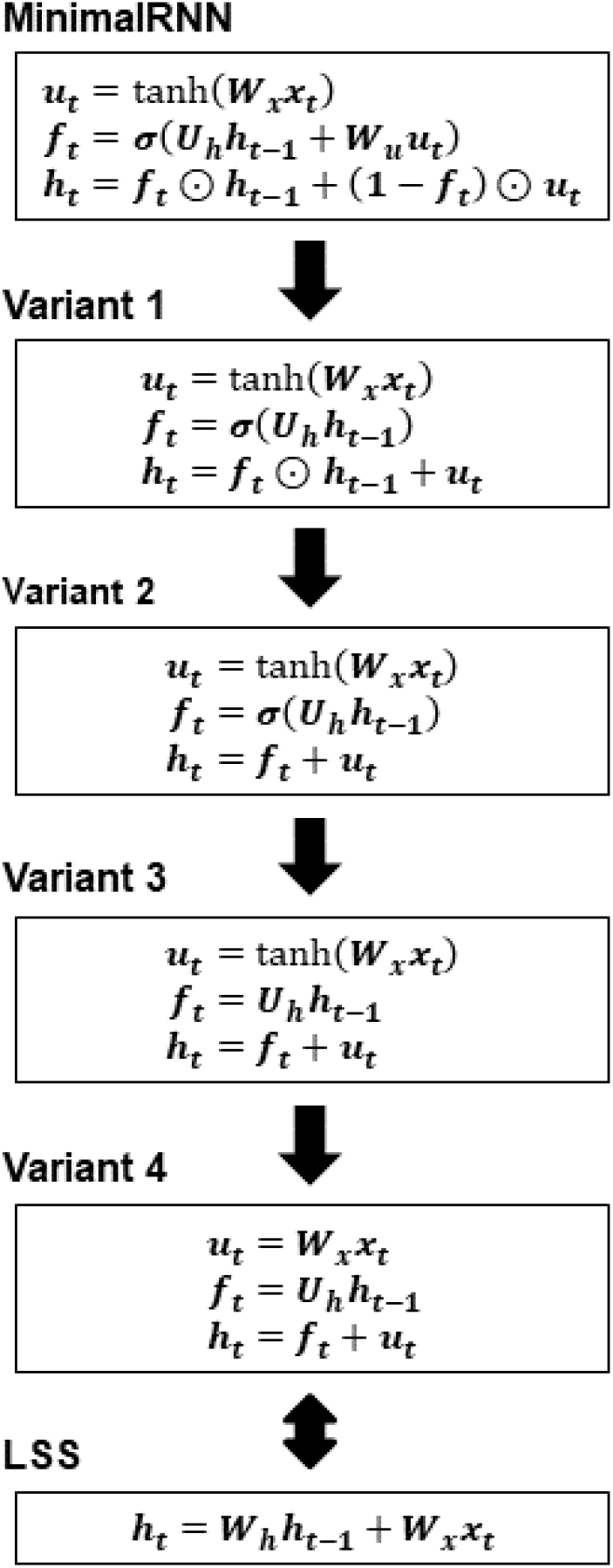
Different ablated minimalRNN models. Ablation is done by simplifying the update equations.

#### 2.7.4 Impact of different features on prediction performance

We performed feature ablation to analyze the contributions of different features to prediction performance of the trained minimalRNN model. To ablate a feature in the input data, the value of that feature was set to the mean value in the dataset, while the other input features were left unaltered. Thus, there were 23 different versions of input data, whereby each version has a different feature ablated. We used the trained minimalRNN model from each split of the data (as described in Section 2.6) and the ablated input data to make prediction in the test data. A large drop in prediction performance when a feature was ablated would suggest that the feature was important for the trained minimalRNN model to make accurate predictions.

### 2.8 TADPOLE live leaderboard

The TADPOLE challenge involves the prediction of ADAS-Cog13, ventricular volume and clinical diagnosis of 219 ADNI participants for every month up to five years into the future. We note that these 219 participants were a subset of the 1677 subjects used in this study. However, the future timepoints used to evaluate performance on the live leaderboard (https://tadpole.grand-challenge.org/D4_Leaderboard/) were not part of the data utilized in this study. Here, we utilized the entire dataset (1677 participants) to tune a set of hyperparameters (using HORD) that maximized performance either (1) one year into the future or (2) all years into the future. We then submitted the predictions of the 219 participants to the TADPOLE leaderboard.

### 2.9 Data and code availability

The code used in this paper can be found at https://github.com/ThomasYeoLab/CBIG/tree/master/stable_projects/predict_phenotypes/Nguyen2020_RNNAD. This study utilized data from the publicly available ADNI database (http://adni.loni.usc.edu/data-samples/access-data/). The particular set of participants and features we used is available at the TADPOLE website (https://tadpole.grand-challenge.org/).

## 3 Results

### 3.1 Overall performance

Figure 9 illustrates the test performance of minimalRNN and four baselines (LSS, LSTM, constant prediction, and SVM/SVR). For brevity, we denote minimalRNN as RNN in all subsequent figures and tables. For clarity, we only showed minimalRNN with model filling (RNN–MF), LSS with model filling (LSS–MF), LSTM with model filling (LSTM-MF) and SVM/SVR using one input timepoint because they yielded the best results within their model classes. Table 4 shows the test performance of all models across all three missing data strategies.

**Figure 9.**
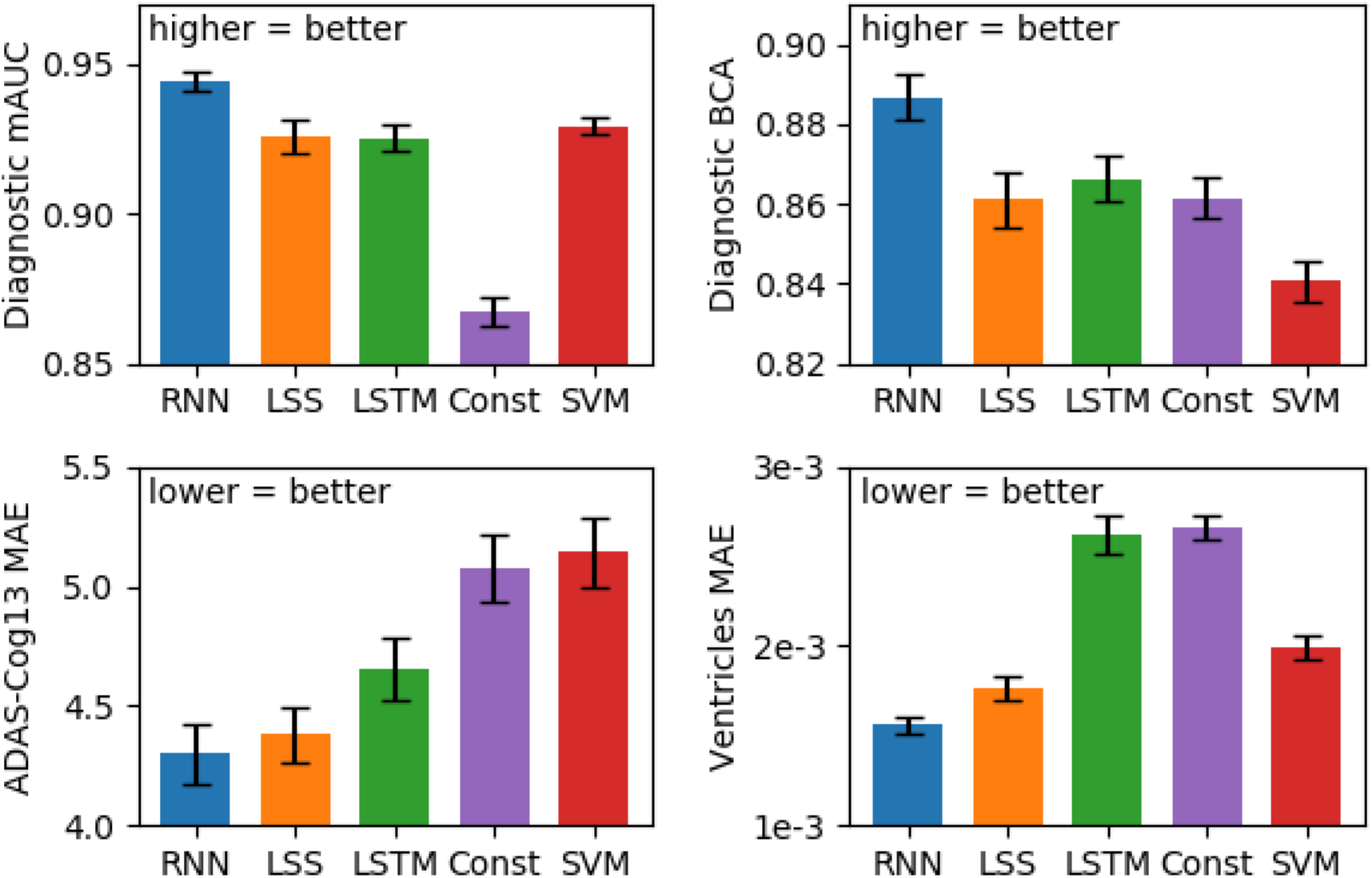
Performance of the best models from each model class averaged across 20 test sets. Error bars show standard error across test sets. For clinical diagnosis, higher mAUC and BCA values indicate better performance. For ADAS-Cog13 and ventricles, lower MAE indicates better performance. For brevity, we denote minimalRNN as RNN. The RNN, LSS, LSTM, and SVM/SVR models corresponded to RNN–MF, LSS-MF, LSTM-MF, and SVM/SVR (= 1tp) in Table 4 respectively. MinimalRNN performed the best. See Figure S1 for all models.

**Table 4.** Prediction performance averaged across 20 test sets. For clinical diagnosis, higher mAUC and BCA values indicate better performance. For ADAS-Cog13 and Ventricles, lower MAE indicates better performance. FF indicates forward filling. LF indicates linear filling. MF indicates model filling. SVM/SVR (= 1tp) utilized one input timepoint. SVM/SVR (≤ 2tp) utilized at most 2 input timepoints (see Section 2.5.2 for details) and so on. The best result for each performance metric was bolded. RNN–MF was numerically the best across all metrics. For brevity, we denote minimalRNN as RNN. Statistical tests were performed between all three minimalRNN variants (RNN-FF, RNN-LF, RNN-MF) and all baseline approaches. Multiple comparisons were corrected using a false discovery rate (FDR) of q < 0.05. Only p-values for RNN-MF are shown. Normal font indicates that RNN-MF was statistically better, while gray font indicates that RNN-MF was not statistically better after FDR correction. The results of SVM/SVR with MFPCA filling are shown in Table S14.

|  | mAUC<br>(more=better) | BCA<br>(more=better) | ADAS-Cog13<br>(less=better) | Ventricles<br>(less=better) |
| --- | --- | --- | --- | --- |
| RNN–FF | 0.923 ± 0.019 | 0.867 ± 0.023 | 5.03 ± 0.62 | 0.00247 ± 0.00036 |
| RNN–LF | 0.910 ± 0.031 | 0.858 ± 0.028 | 5.42 ± 0.94 | 0.00193 ± 0.00029 |
| RNN–MF | **0.944** ± 0.014 | **0.887** ± 0.024 | **4.30** ± 0.53 | **0.00156** ± 0.00022 |
| LSS–FF | 0.928 ± 0.020<br>(p = 0.018) | 0.864 ± 0.024<br>(p = 0.001) | 4.95 ± 0.57<br>(p = 0.003) | 0.00216 ± 0.00031<br>(p = 5.6×10<sup>-7</sup>) |
| LSS–LF | 0.908 ± 0.032<br>(p = 0.005) | 0.857 ± 0.037<br>(p = 0.042) | 6.36 ± 0.82<br>(p = 3.2×10<sup>-7</sup>) | 0.00175 ± 0.00023<br>(p = 0.061) |
| LSS–MF | 0.926 ± 0.025<br>(p = 0.004) | 0.861 ± 0.029<br>(p = 0.001) | 4.38 ± 0.49<br>(p = 0.590) | 0.00177 ± 0.00028<br>(p = 0.044) |
| LSTM–FF | 0.932 ± 0.018<br>(p = 0.033) | 0.857 ± 0.029<br>(p = 1.4e-03) | 5.13 ± 0.58<br>(p = 3.6e-04) | 0.00360 ± 0.00087<br>(p = 2.7e-06) |
| LSTM–LF | 0.920 ± 0.021<br>(p = 1.8e-04) | 0.871 ± 0.028<br>(p = 0.031) | 5.38 ± 0.73<br>(p = 1.4e-04) | 0.00477 ± 0.00065<br>(p = 3.4e-11) |
| LSTM–MF | 0.925 ± 0.019<br>(p = 1.8e-03) | 0.866 ± 0.025<br>(p = 4.3e-04) | 4.65 ± 0.56<br>(p = 0.031) | 0.00263 ± 0.00047<br>(p = 3.8e-06) |
| Constant | 0.867 ± 0.022<br>(p = 3.2×10<sup>-9</sup>) | 0.861 ± 0.023<br>(p = 2.0×10<sup>-4</sup>) | 5.07 ± 0.61<br>(p = 3.3×10<sup>-4</sup>) | 0.00266 ± 0.00027<br>(p = 5.9×10<sup>-12</sup>) |
| SVM/SVR (= 1tp) | 0.929 ± 0.013<br>(p = 0.011) | 0.841 ± 0.023<br>(p = 2.5×10<sup>-7</sup>) | 5.14 ± 0.62<br>(p = 1.8×10<sup>-4</sup>) | 0.00199 ± 0.00031<br>(p = 7.3×10<sup>-5</sup>) |
| SVM/SVR (≤ 2tp) | 0.926 ± 0.013<br>(p = 0.002) | 0.836 ± 0.026<br>(p = 2.8×10<sup>-6</sup>) | 5.23 ± 0.63<br>(p = 1.1×10<sup>-4</sup>) | 0.00230 ± 0.00037<br>(p = 2.7×10<sup>-7</sup>) |
| SVM/SVR (≤ 3tp) | 0.923 ± 0.013<br>(p = 0.001) | 0.830 ± 0.025<br>(p = 2.6×10<sup>-7</sup>) | 5.53 ± 0.55<br>(p = 4.5×10<sup>-7</sup>) | 0.00261 ± 0.00037<br>(p = 5.9×10<sup>-7</sup>) |
| SVM/SVR (≤ 4tp) | 0.919 ± 0.012<br>(p = 2.2×10<sup>-5</sup>) | 0.832 ± 0.019<br>(p = 4.1×10<sup>-7</sup>) | 5.68 ± 0.58<br>(p = 9.4×10<sup>-7</sup>) | 0.00269 ± 0.00035<br>(p = 1.2×10<sup>-9</sup>) |

We performed statistical tests comparing the three minimalRNN variants (RNN–FF, RNN–LF and RNN–MF) with all other baseline approaches (LSS, LSTM, constant prediction, SVM/SVR). Multiple comparisons were corrected with a false discovery rate (FDR) of q < 0.05. RNN-MF showed the best results and was statistically better than most baseline approaches (Table 4). For example, RNN-MF was statistically better than LSS-MF for clinical diagnosis, but not ADAS-Cog13 or ventricular volume. Similarly, RNN-MF was statistically better than LSTM-MF for clinical diagnosis and ventricular volume, but not ADAS-Cog13.

In terms of handling missing data, model filling (MF) performed better than forward filling (FF) and linear filling (LF), especially when predicting ADAS-Cog13 and ventricular volume (Table 4). Interestingly, more input timepoints do not necessarily lead to better prediction in the case of SVM/SVR. In fact, the SVM/SVR model using one timepoint was numerically better than SVM/SVR models using more timepoints, although differences were small. This might be because SVM/SVR models with one input timepoint had access to more training data than SVM/SVR models with more input timepoints (Section 2.5.2). Furthermore, SVM/SVR models with more input timepoints had to handle longer feature vectors, which increased the risk of overfitting (Section 2.5.2).

Recall that for test subjects, the first half of the timepoints of each subject were used to predict the second half of the timepoints of the same subject (Section 2.6). Table 5 shows the breakdown of subjects based on their clinical diagnoses at the last input timepoints (with observed clinical diagnoses) and the last timepoints (with observed clinical diagnoses). For example, if a subject had 10 timepoints, then the 10 timepoints were split into 5 input (observed) timepoints and 5 unobserved timepoints we seek to predict. Then, in the case of this subject, the last input timepoint would be timepoint 5 and the last timepoint would be timepoint 10. If the subject did not have observed clinical diagnosis at timepoint 10, then we would consider the clinical diagnosis at timepoint 9 and so on. We note that a small number of subjects was not included in Table 5 because they did not have any observed clinical diagnosis in the first half and/or second half of the timepoints.

**Table 5.**
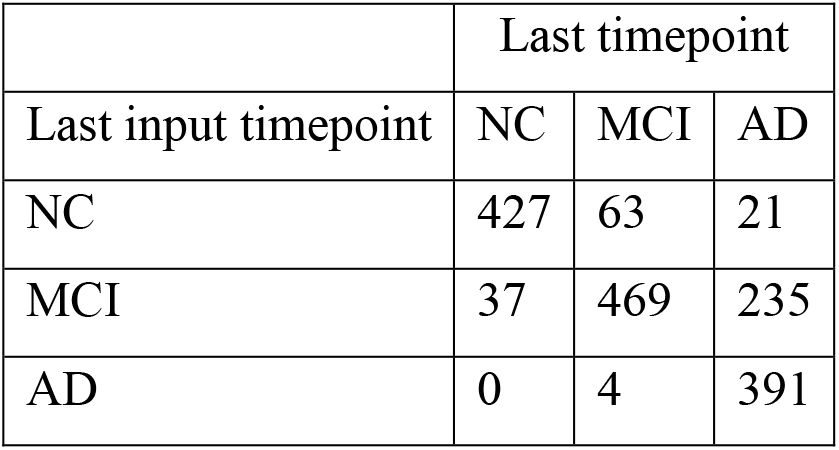
Breakdown of subjects based on their clinical diagnoses at the last input timepoints (with observed clinical diagnoses) and the last timepoints (with observed clinical diagnoses).

Figure 10 shows the breakdown of the prediction performance (Figure 9) into six different groups. The “stable” groups (NC-S, MCI-S, AD) comprised subjects whose diagnostic categories were the same at the last input timepoint and the last timepoint. The “progressive” groups (NC-P, MCI-P) comprised subjects who progressed along the AD dementia spectrum (e.g., from MCI to AD). Finally, the MCI recovered (MCI-R) group comprised subjects who have reverted from MCI to NC. We did not consider the 4 subjects that reverted from AD to MCI because of the small sample size. We note that diagnostic prediction performance was measured using accuracy (fraction of correct predictions) instead of mAUC and BCA because there was only one class in the stable groups.

**Figure 10.**
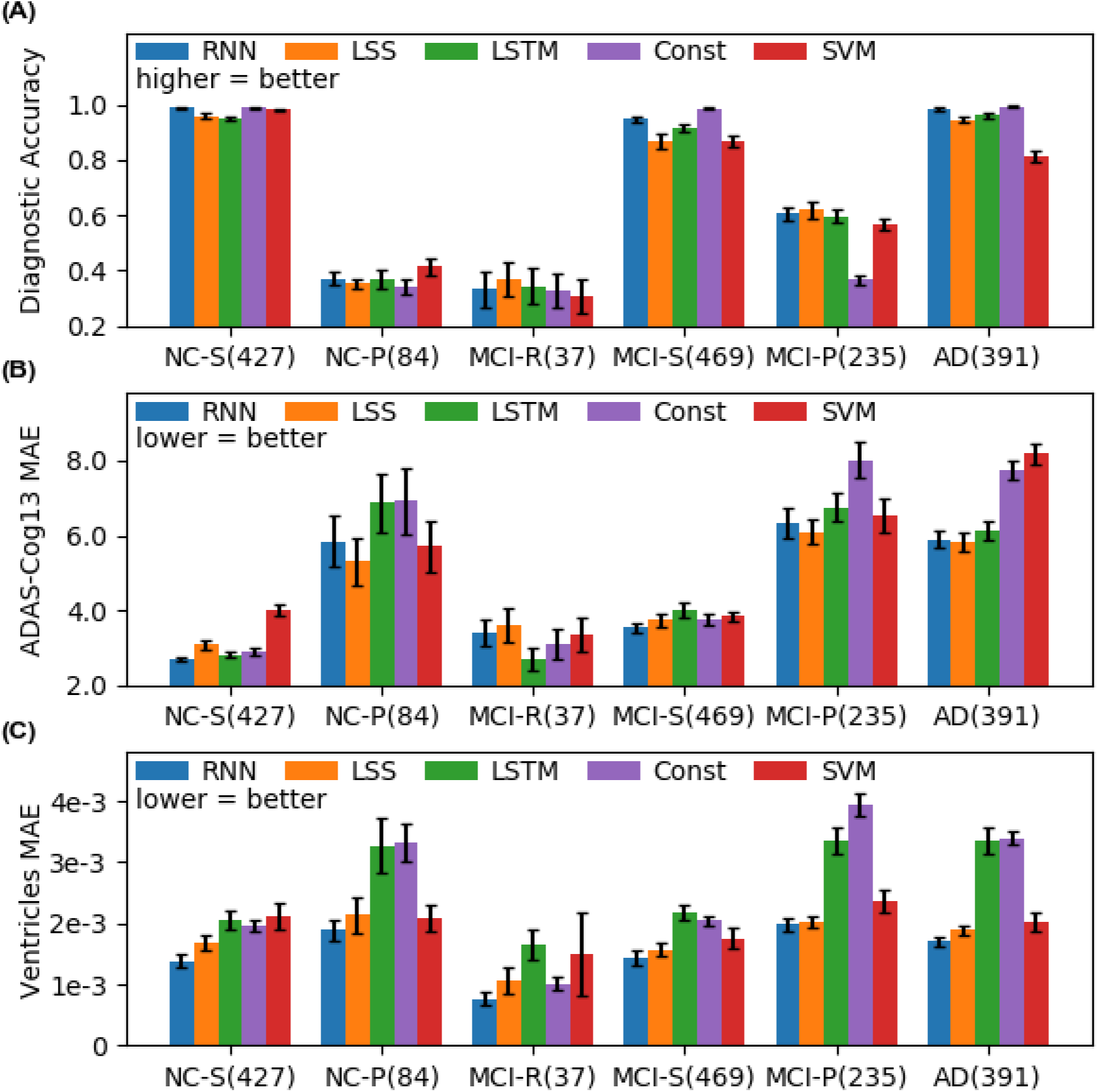
Prediction performance broken down into six different groups: NC stable (NC-S), NC progressive (NC-P), MCI recovered (MCI-R), MCI stable (MCI-stable), MCI progressive (MCI-P) and AD stable (AD). The numbers in the brackets indicate the numbers of subjects in the respective groups. For brevity, we denote minimalRNN as RNN. The minimalRNN compared favorably with all baseline algorithms in almost all groups.

In the case of predicting ventricular volume or ADAS-Cog13, minimalRNN was comparable to or numerically better than all baselines. In the case of diagnostic category, minimalRNN compared favorably with all baselines except for constant prediction in the stable groups. The reason is that it is optimal to predict all future diagnostic categories to be the same as the last observed diagnosis in the stable groups. However, in reality, whether subjects are stable or not is not known in advance. Therefore, for the stable groups, constant prediction should be treated as an upper bound on prediction performance, rather than a baseline. We note constant prediction did not achieve 100% accuracy in the stable groups because the clinical diagnoses could fluctuate over time. For example, if a subject had 4 timepoints with corresponding diagnoses NC, NC, MCI and NC. Then, the subject would be classified as NC-stable because the second and fourth timepoints had the same NC diagnoses.

Figure 11 shows the breakdown of the prediction performance from Figure 9 in yearly interval up to 6 years into the future. Not surprisingly, the performance of all algorithms became worse for predictions further into the future. The constant baseline was very competitive against the other models for the first year, but performance for subsequent years dropped very quickly. The minimalRNN model was comparable or numerically better than all baseline approaches across all the years.

**Figure 11.**
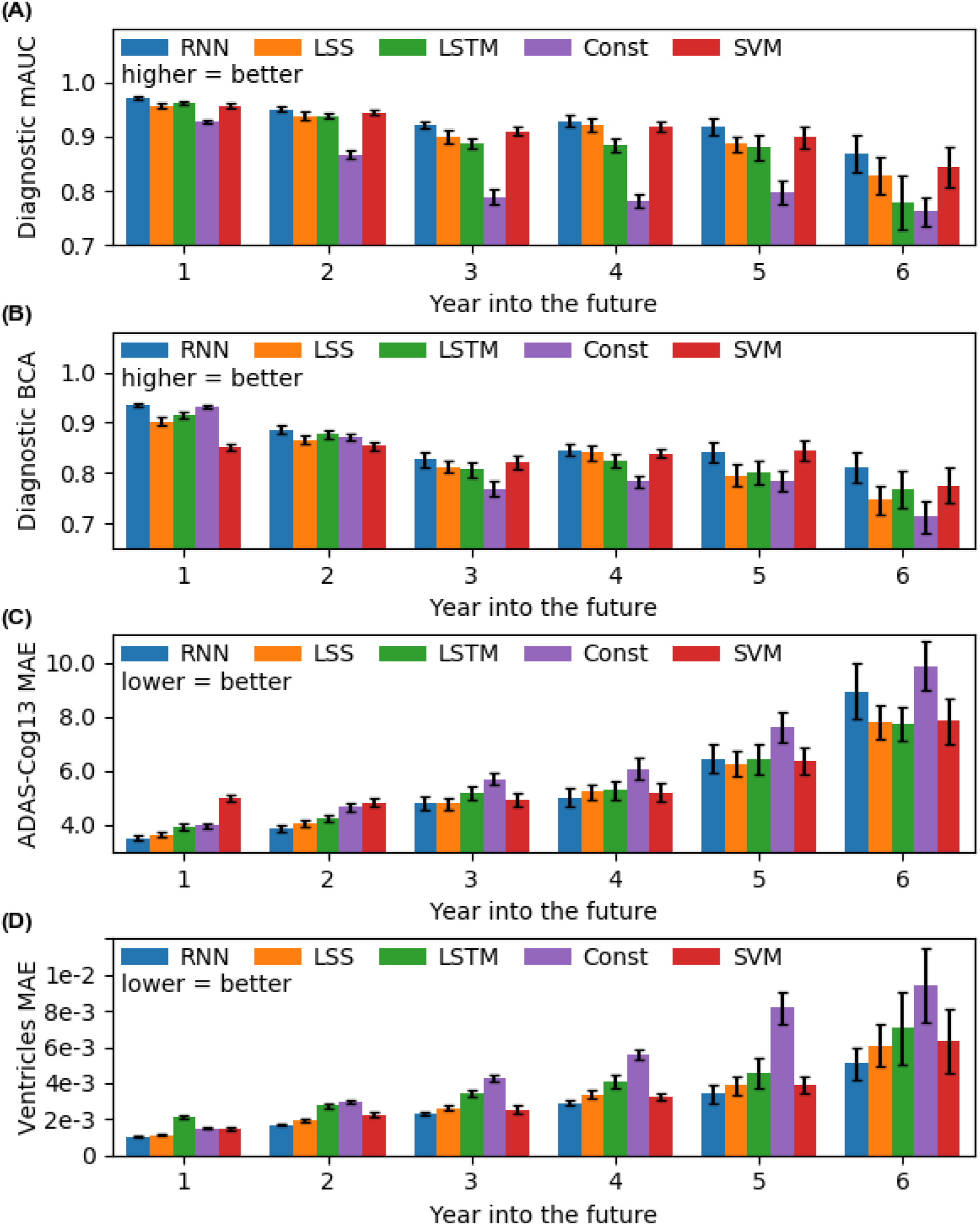
Prediction performance from Figure 9 broken down into yearly interval up to 6 years into the future. For brevity, we denote minimalRNN as RNN. All algorithms became worse further into the future. MinimalRNN was comparable to or numerically better than all baseline algorithms across all years. See Figure S2 for all models.

### 3.2 Further analysis

#### 3.2.1 MinimalRNN using one and four input timepoints in test subjects achieve comparable performance

Given that the MinimalRNN with model filling (RNN–MF) performed the best (Table 4), we further explored how well the trained RNN–MF model would perform on test subjects with different number of input timepoints. Figure 12 shows the performance of RNN-MF averaged across 20 test sets using different number of input timepoints. The exact numerical values are reported in Table 6. RNNs using 2 to 4 input timepoints achieved similar performance across all metrics. RNN using 1 input timepoint had numerically worse results, especially for ventricular volume. However, there was no statistical difference between using 1 input timepoint and 4 input timepoints even in the case of ventricular volume (p = 0.20).

**Figure 12.**
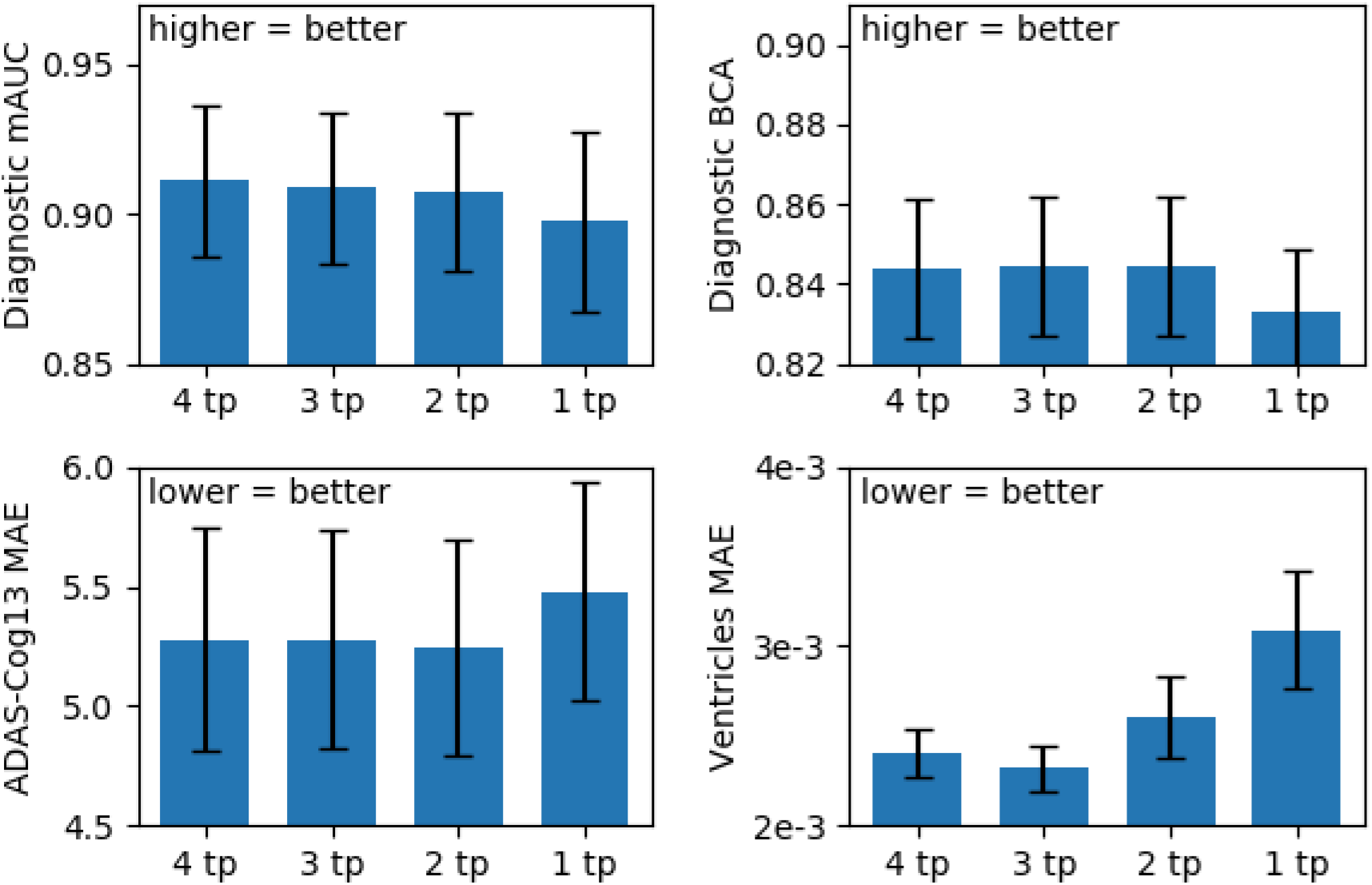
Test performance of minimalRNN model with model filling strategy (RNN-MF) using different numbers of input timepoints (after training with all timepoints). Results were averaged across 20 test sets. Even though the minimalRNN model using 1 input timepoint yielded numerically worse results, the differences were not significant (see Table 6).

**Table 6.** Test performance of minimalRNN model with model filling strategy (RNN-MF) using different numbers of input timepoints (after training with all timepoints). Results were averaged across 20 test sets. Statistical tests were performed to test for differences between using 4 timepoints versus less timepoints. The gray font indicates that there was no statistical difference that survived FDR of q < 0.05.

|  | mAUC<br>(more=better) | BCA<br>(more=better) | ADAS-Cog13<br>(less=better) | Ventricles<br>(less=better)s |
| --- | --- | --- | --- | --- |
| 4 timepoints | 0.911 ± 0.076 | 0.844 ± 0.053 | 5.28 ± 1.41 | 0.00240 ± 0.00040 |
| 3 timepoints | 0.909 ± 0.076<br>(p = 0.68) | 0.844 ± 0.052<br>(p = 0.88) | 5.28 ± 1.38<br>(p = 0.99) | 0.00232 ± 0.00038<br>(p = 0.22) |
| 2 timepoints | 0.908 ± 0.080<br>(p = 0.57) | 0.844 ± 0.053<br>(p = 0.84) | 5.24 ± 1.35<br>(p = 0.89) | 0.00260 ± 0.00067<br>(p = 0.50) |
| 1 timepoint | 0.897 ± 0.091<br>(p = 0.27) | 0.833 ± 0.048<br>(p = 0.18) | 5.48 ± 1.37<br>(p = 0.53) | 0.00309 ± 0.00098<br>(p = 0.20) |

#### 3.2.2 Varying temporal resolution has little impact on performance

Table 7 shows the prediction performance of the RNN-MF model when the temporal resolution varied from 1-month interval to 6-month interval. There was no significant difference in prediction performance across different temporal resolutions.

**Table 7.** Test performance of minimalRNN model with model filling strategy (RNN-MF) at different temporal resolution. We note that the top row (1-month interval) was the same as in Table 4. Results were averaged across 20 test sets. The best result for each performance metric was bolded. There was no significant difference across different temporal resolutions.

|  | mAUC<br>(more=better) | BCA<br>(more=better) | ADAS-Cog13<br>(less=better) | Ventricles<br>(less=better) |
| --- | --- | --- | --- | --- |
| 1-month interval | **0.944** ± 0.014 | **0.887** ± 0.024 | 4.30 ± 0.53 | 0.00156 ± 0.00022 |
| 3-month interval | 0.942 ± 0.016<br>(p = 0.58) | 0.886 ± 0.026<br>(p = 0.88) | **4.11** ± 0.49<br>(p = 0.079) | **0.00153** ± 0.00014<br>(p = 0.59) |
| 6 month interval | 0.940 ± 0.017<br>(p = 0.27) | 0.885 ± 0.023<br>(p = 0.75) | 4.13 ± 0.51<br>(p = 0.22) | 0.00158 ± 0.00021<br>(p = 0.79) |

#### 3.2.3 Impact of different terms in the minimalRNN model

Table 8 shows the performances of the original minimalRNN model (RNN-MF) and 4 ablated variants decreasing in complexity from RNN-MF to variant 4 (LSS-MF). Numerically, RNN-MF had the best results compared with all 4 variants. However, it was not the case that performance continually degraded from the most complex model (RNN-MF) to the least complex model (LSS-MF). Interestingly, among the 4 variants, LSS-MF (Variant 4) showed the worst performance for clinical diagnosis, but close to the best performance for ADAS-Cog13 and ventricular volume. This suggests that some level of nonlinearity might be more useful for predicting clinical diagnosis, but less so for ADAS-Cog13 and ventricular volume. Overall, it was difficult to conclude that a specific component was essential to minimalRNN’s performance. This might not be surprising because as its name suggested, the minimalRNN was designed to be as simple as possible, so removing any component yielded somewhat worse results.

**Table 8.** Test performance of the original minimalRNN model (RNN-MF) and different ablated variants. Results were averaged across 20 test sets. The best result for each performance metric was **bolded.**

|  | mAUC<br>(more=better) | BCA<br>(more=better) | ADAS-Cog13<br>(less=better) | Ventricles<br>(less=better) |
| --- | --- | --- | --- | --- |
| RNN-MF | **0.944 ± 0.014** | **0.887 ± 0.024** | **4.30 ± 0.53** | **0.00156 ± 0.00022** |
| Variant 1 | 0.934 ± 0.018 | 0.878 ± 0.022 | 4.59 ± 0.53 | 0.00200 ± 0.00055 |
| Variant 2 | 0.928 ± 0.019 | 0.868 ± 0.034 | 4.41 ± 0.43 | 0.00179 ± 0.00040 |
| Variant 3 | 0.932 ± 0.013 | 0.876 ± 0.021 | 4.32 ± 0.49 | 0.00186 ± 0.00034 |
| Variant 4<br>(LSS-MF) | 0.926 ± 0.025 | 0.861 ± 0.029 | 4.38 ± 0.49 | 0.00177 ± 0.00028 |

#### 3.2.4 Impact of different features on prediction performance

The results of the feature ablation experiments are shown in Table 9. Unsurprisingly, ablating diagnosis resulted in the most significant drop in diagnostic mAUC and BCA, while ablating ADAS-Cog13 and ventricular volume resulted in the most significant increase in ADAS-Cog13 MAE and ventricular MAE respectively. Ablating CDRSB also led to a noticeable drop in diagnosis mAUC and BCA, probably because CDRSB is used in the diagnosis of an individual. Interestingly, ablating CDRSB also led to a noticeable increase in ventricular MAE.

**Table 9.** Test performance of minimalRNN model (RNN-MF) with different features ablated (replacing input feature with the mean value). Results were averaged across 20 test sets. Prediction performance of the original model was bolded. For each column, the top two ablated features leading to the largest drop in performance were bolded and italicized.

|  | mAUC<br>(more=better) | BCA<br>(more=better) | ADAS-Cog13<br>(less=better) | Ventricles<br>(less=better) |
| --- | --- | --- | --- | --- |
| No Ablation | **0.944 ± 0.014** | **0.887 ± 0.024** | **4.30 ± 0.53** | **0.00156 ± 0.00022** |
| Ablate CDRSB | **0.916 ± 0.049** | **0.858 ± 0.055** | 4.29 ± 0.48 | **0.00162 ± 0.00025** |
| Ablate ADAS-Cog11 | 0.943 ± 0.015 | 0.884 ± 0.024 | **5.15 ± 0.95** | 0.00161 ± 0.00022 |
| Ablate ADAS-Cog13 | 0.941 ± 0.021 | 0.875 ± 0.029 | **6.96 ± 3.31** | 0.00160 ± 0.00025 |
| Ablate MMSE | 0.945 ± 0.014 | 0.882 ± 0.023 | 4.43 ± 0.56 | 0.00157 ± 0.00021 |
| Ablate RAVLT<br>immediate | 0.942 ± 0.016 | 0.882 ± 0.025 | 4.69 ± 0.62 | 0.00155 ± 0.00022 |
| Ablate RAVLT<br>learning | 0.943 ± 0.014 | 0.884 ± 0.023 | 4.33 ± 0.52 | 0.00159 ± 0.00022 |
| Ablate RAVLT<br>forgetting | 0.945 ± 0.015 | 0.887 ± 0.023 | 4.29 ± 0.52 | 0.00155 ± 0.00021 |
| Ablate RAVLT<br>forgetting percent | 0.935 ± 0.028 | 0.878 ± 0.029 | 4.89 ± 1.19 | 0.00165 ± 0.00024 |
| Ablate Functional<br>Activities<br>Questionnaire (FAQ) | 0.943 ± 0.016 | 0.882 ± 0.026 | 4.29 ± 0.45 | 0.00155 ± 0.00020 |
| Ablate Montreal<br>Cognitive Assessment<br>(MOCA) | 0.944 ± 0.015 | 0.883 ± 0.026 | 4.56 ± 0.59 | 0.00155 ± 0.00021 |
| Ablate Ventricles | 0.944 ± 0.014 | 0.887 ± 0.025 | 4.29 ± 0.49 | **0.00166 ± 0.00017** |
| Ablate Hippocampus | 0.941 ± 0.014 | 0.884 ± 0.025 | 4.40 ± 0.58 | 0.00158 ± 0.00021 |
| Ablate Whole brain<br>volume | 0.945 ± 0.015 | 0.886 ± 0.024 | 4.30 ± 0.53 | 0.00157 ± 0.00021 |
| Ablate Entorhinal<br>cortical volume | 0.944 ± 0.015 | 0.883 ± 0.025 | 4.33 ± 0.55 | 0.00156 ± 0.00021 |
| Ablate Fusiform<br>cortical volume | 0.944 ± 0.014 | 0.883 ± 0.024 | 4.29 ± 0.50 | 0.00156 ± 0.00022 |
| Ablate Middle temporal<br>cortical volume | 0.945 ± 0.015 | 0.884 ± 0.024 | 4.33 ± 0.50 | 0.00156 ± 0.00022 |
| Ablate Intracranial<br>volume | 0.945 ± 0.014 | 0.886 ± 0.025 | 4.29 ± 0.53 | 0.00156 ± 0.00020 |
| Ablate Florbetapir<br>(18F-AV-45) - PET | 0.944 ± 0.015 | 0.887 ± 0.024 | 4.29 ± 0.52 | 0.00155 ± 0.00020 |
| Ablate<br>Fluorodeoxyglucose<br>(FDG) - PET | 0.943 ± 0.014 | 0.883 ± 0.025 | 4.30 ± 0.54 | 0.00155 ± 0.00021 |
| Ablate Beta-amyloid<br>(CSF) | 0.944 ± 0.016 | 0.884 ± 0.025 | 4.33 ± 0.51 | 0.00156 ± 0.00022 |
| Ablate Total tau | 0.944 ± 0.015 | 0.885 ± 0.025 | 4.34 ± 0.54 | 0.00156 ± 0.00021 |
| Ablate Phosphorylated<br>tau | 0.943 ± 0.014 | 0.885 ± 0.023 | 4.37 ± 0.55 | 0.00156 ± 0.00021 |
| Ablate Diagnosis | **0.878 ± 0.032** | **0.770 ± 0.031** | 4.31 ± 0.43 | 0.00157 ± 0.00021 |

### 3.3 TADPOLE live leaderboard

The original LSTM model (Nguyen et al., 2018) was ranked 5th (out of 53 entries) in the TADPOLE grand challenge in July 2019 (entry “CBIL” in https://tadpole.grand-challenge.org/Results/). Our current minimalRNN models were ranked 2nd and 3rd (out of 63 entries) on the leaderboard as of June 3rd, 2020 (entries (“CBIL-MinMFa” and “CBIL-MinMF1”; https://tadpole.grand-challenge.org/D4_Leaderboard/). Interestingly, the model obtained from hyperparameters tuned to predict all years into the future (“CBIL-MinMFa”) performed better than the model obtained from hyperparameters tuned to predict one year into the future (“CBIL-MinMF1”), even though the leaderboard currently utilized about one year of future data for prediction.

## 4 Discussion

In this work, we adapted a minimalRNN model for predicting longitudinal progression in AD dementia. Our approach compared favorably with baseline algorithms, such as SVM/SVR, LSS, and LSTM models. However, we note that there was no statistical difference between the minimalRNN and LSS for predicting ADAS-Cog13 and ventricular volume even though other studies suggested benefits of modeling non-linear interactions between features (Popescu et al., 2019).

As can be seen when setting up the SVM/SVR baseline models (Section 2.5.2), there were a lot of edge cases to consider in order to adapt a “static” prediction algorithm (e.g., SVM/SVR) to the more “dynamic” longitudinal prediction problem we considered here. For example, data is wasted because static approaches generally assume that participants have the same number of input timepoints. Therefore, for the SVM/SVR models using 4 input timepoints, we ended up with only 1454 participants out of the original 1677 participants. This might explain why the SVM/SVR model using 1 input timepoint compared favorably with the SVM/SVR model using 4 input timepoints (Table 4). Furthermore, we had to build multiple separate SVM/SVR models to predict at a fixed number of future timepoints, and performed interpolation at intermediate timepoints. By contrast, state-based models (e.g., minimalRNN, LSS, or LSTM) are more elegant in the sense that they handle participants with different number of timepoints and can in principle predict unlimited number of timepoints into the future.

Even though the ADNI dataset comprised participants with multiple timepoints, for the algorithm to be clinically useful, it has to be successful at dealing with missing data and participants with only one input timepoint. We found that the “integrative” approach of using the model to fill in the missing data (i.e., model filling) compared favorably with “preprocessing” approaches, such as forward filling or linear filling. However, it is possible that more sophisticated “preprocessing” approaches, such as matrix factorization (Mazumder et al., 2010; Nie et al., 2017; Thung et al., 2016) or wavelet interpolation (Mondal and Percival, 2010), might yield better results. We note that our model filling approach can also be considered as a form of matrix completion since the RNN (or LSS) was trained to minimize the predictive loss, which is equivalent to maximizing the likelihood of the training data. However, matrix completion usually assumes that the training data can be represented as a matrix that can be factorized into low-ranked or other specially-structured matrices. On the other hand, our method assumes temporal dependencies between rows in the data matrix (where each row is a timepoint).

Our best model (minimalRNN with model filling) had similar performance when using only 1 input timepoint instead of 4 input timepoints, suggesting that our approach might work well with just cross-sectional data (after training using longitudinal data). However, we might have simply lacked the statistical power to distinguish among the different conditions because of the smaller number of subjects in this experiment. Overall, there was no noticeable difference among using 2, 3 or 4 input timepoints, while the performance using 1 input timepoint appeared worse, but the difference was not statistically significant (Figure 12).

Although our approach compared favorably with the baseline algorithms, we note that any effective AD dementia treatment probably has to begin early in the disease process, potentially at least a decade before the emergence of behavioral symptoms. However, even in the case of our best model (minimalRNN with model filling), prediction performance of clinical diagnosis dropped from a BCA of 0.935 in year 1 to a BCA of 0.810 in year 6, while ventricular volume MAE increased from 0.00104 in year 1 to 0.00511 in year 6. Thus, significant improvement is needed for clinical utility.

One possible future direction is to investigate new features, e.g., those derived from diffusion MRI or arterial spin labeling. Previous studies have also suggested that different atrophy patterns (beyond the temporal lobe) might influence cognitive decline early in the disease process (Noh et al., 2014; Byun et al., 2015; Ferreira et al., 2017; Zhang et al., 2016; Risacher et al., 2017; Sun et al., 2019), so the atrophy features considered in this study (Table 1) might not be optimal. Although the new features may be correlated with currently used features, the new features might still provide complementary information when modeling AD progression (Popescu et al., 2019).

As mentioned in the introduction, an earlier version of our algorithm was ranked 5th out of 50 entries in the TADPOLE competition. Our current model was ranked 2nd out of 63 entries on the TADPOLE live leaderboard as of June 2nd, 2020. Interestingly, the top team considered additional handcrafted features, which might have contributed to its success. Furthermore, the top team utilized a non-deep-learning algorithm XGboost (Chen and Guestrin, 2016), which might be consistent with recent work suggesting that for certain neuroimaging applications, non-deep-learning approaches might be highly competitive (He et al., 2020)

## 5 Conclusion

Using 1677 participants from the ADNI database, we showed that the minimalRNN model was better than other baseline algorithms for the longitudinal prediction of multimodal AD biomarkers and clinical diagnosis of participants up to 6 years into the future. We explored three different strategies to handle the missing data issue prevalent in longitudinal data. We found that the RNN model can itself be used to fill in the missing data, thus providing an integrative strategy to handle the missing data issue. Furthermore, we also found that after training with longitudinal data, the trained RNN model can perform reasonably well using one input timepoint, suggesting the approach might also work for cross-sectional data.

## Acknowledgment

This work was supported by the Singapore National Research Foundation (NRF) Fellowship (Class of 2017). Any opinions, findings and conclusions or recommendations expressed in this material are those of the author(s) and do not reflect the views of National Research Foundation, Singapore. Our research also utilized resources provided by the Center for Functional Neuroimaging Technologies, P41EB015896 and instruments supported by 1S10RR023401, 1S10RR019307, and 1S10RR023043 from the Athinoula A. Martinos Center for Biomedical Imaging at the Massachusetts General Hospital. Our computational work was partially performed on resources of the National Supercomputing Centre, Singapore (https://www.nscc.sg). The Titan Xp used for this research was donated by the NVIDIA Corporation. Data collection and sharing for this project was funded by the Alzheimer’s Disease Neuroimaging Initiative (ADNI) (National Institutes of Health Grant U01 AG024904) and DOD ADNI (Department of Defense award number W81XWH-12-2-0012). ADNI is funded by the National Institute on Aging, the National Institute of Biomedical Imaging and Bioengineering, and through generous contributions from the following: AbbVie, Alzheimer’s Association; Alzheimer’s Drug Discovery Foundation; Araclon Biotech; BioClinica, Inc.; Biogen; Bristol-Myers Squibb Company; CereSpir, Inc.; Cogstate; Eisai Inc.; Elan Pharmaceuticals, Inc.; Eli Lilly and Company; EuroImmun; F. Hoffmann-La Roche Ltd and its affiliated company Genentech, Inc.; Fujirebio; GE Healthcare; IXICO Ltd.; Janssen Alzheimer Immunotherapy Research & Development, LLC.; Johnson & Johnson Pharmaceutical Research & Development LLC.; Lumosity; Lundbeck; Merck & Co., Inc.; Meso Scale Diagnostics, LLC.; NeuroRx Research; Neurotrack Technologies; Novartis Pharmaceuticals Corporation; Pfizer Inc.; Piramal Imaging; Servier; Takeda Pharmaceutical Company; and Transition Therapeutics. The Canadian Institutes of Health Research is providing funds to support ADNI clinical sites in Canada. Private sector contributions are facilitated by the Foundation for the National Institutes of Health (www.fnih.org). The grantee organization is the Northern California Institute for Research and Education, and the study is coordinated by the Alzheimer’s Therapeutic Research Institute at the University of Southern California. ADNI data are disseminated by the Laboratory for Neuro Imaging at the University of Southern California.

## APPENDIX A

This appendix summarizes differences between the minimalRNN and LSTM. For the convenience of the readers, the minimalRNN state equations (Figure 2B) are repeated below.

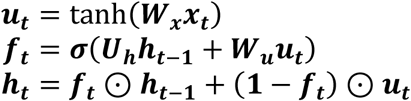

For ease of comparison, we use a similar set of notations to show the LSTM state equations below.

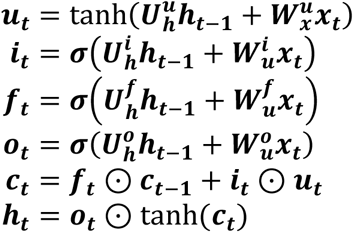

As can be seen, minimalRNN uses fewer parameters than LSTM by doing away with the output gate (***o***_***t***_ ) and setting the input gate ( ***i***_***t***_ ) to be the complement of the forget gate (i.e. (1 − ***f*** _***t***_)). The hyperbolic tangent is also removed from the computation of ***h***_***t***_ , thus making ***h***_***t***_ = ***c***_***t***_. In addition, the term ***h***_***t***−**1**_ is removed from the computation of the term ***u***_***t***_ in the minimalRNN, so the hidden state ( ***h***_***t***_ ) of the minimalRNN decays to zero when the input ( ***x***_***t***_ ) is zero. Note that in the context of our study, all variables (except clinical diagnosis) were z-normalized (Section 2.6). Thus, input of zero corresponds to observing the mean value. In contrast, the hidden state of the LSTM can fluctuate even when the input is zero.

## Supplemental Results

**Figure S1.**
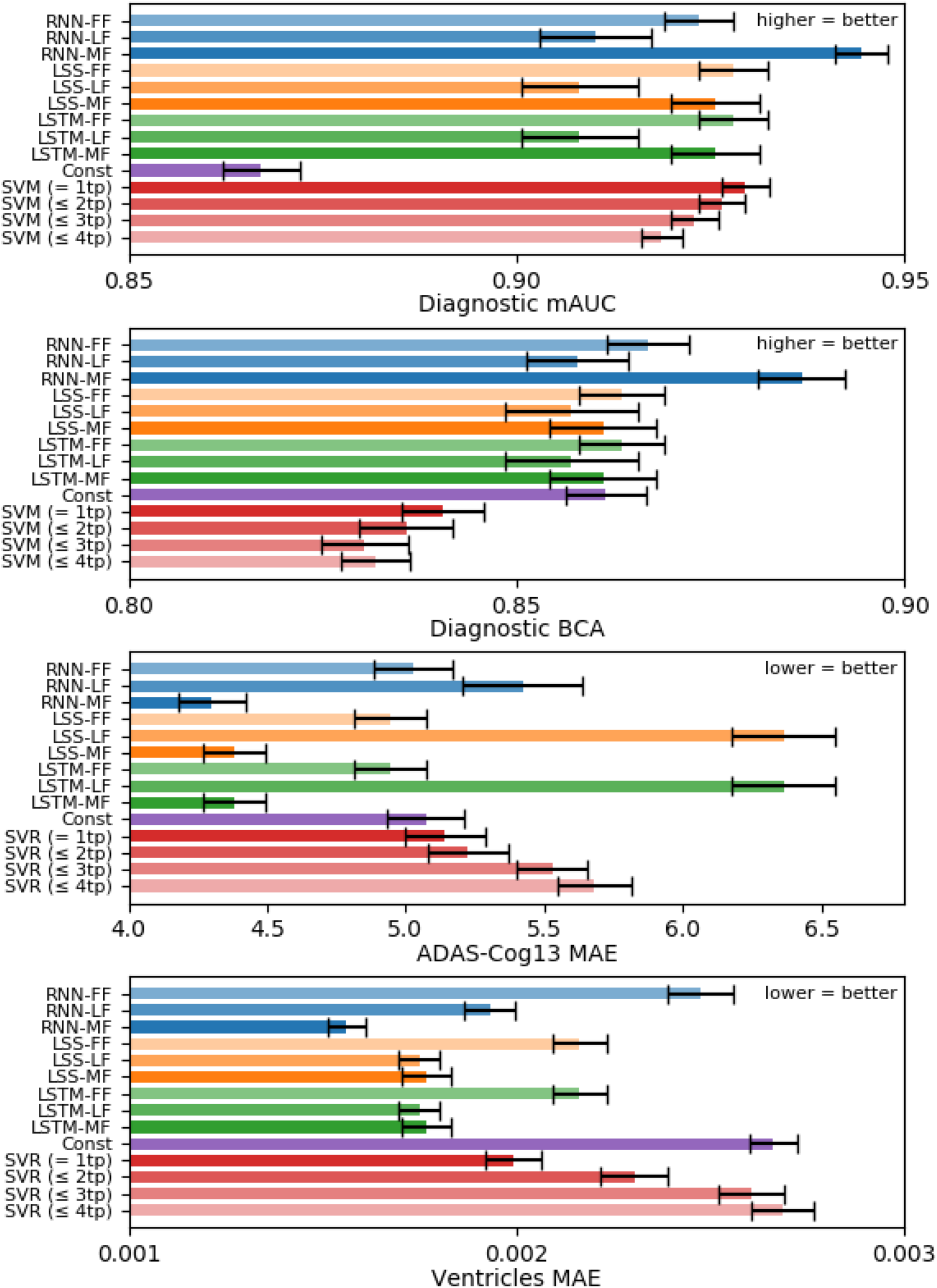
Performance of all models averaged across 20 test sets. Error bars show standard error across test sets. For clinical diagnosis, higher mAUC and BCA values indicate better performance. For ADAS-Cog13 and Ventricles, lower MAE indicates better performance. FF indicates forward filling. LF indicates linear filling. MF indicates model filling. SVM/SVR (= 1tp) utilized one input timepoint. SVM/SVR (≤ 2tp) utilized at most 2 input timepoints (see Section 2.5.2 for details) and so on.

**Figure S2.**
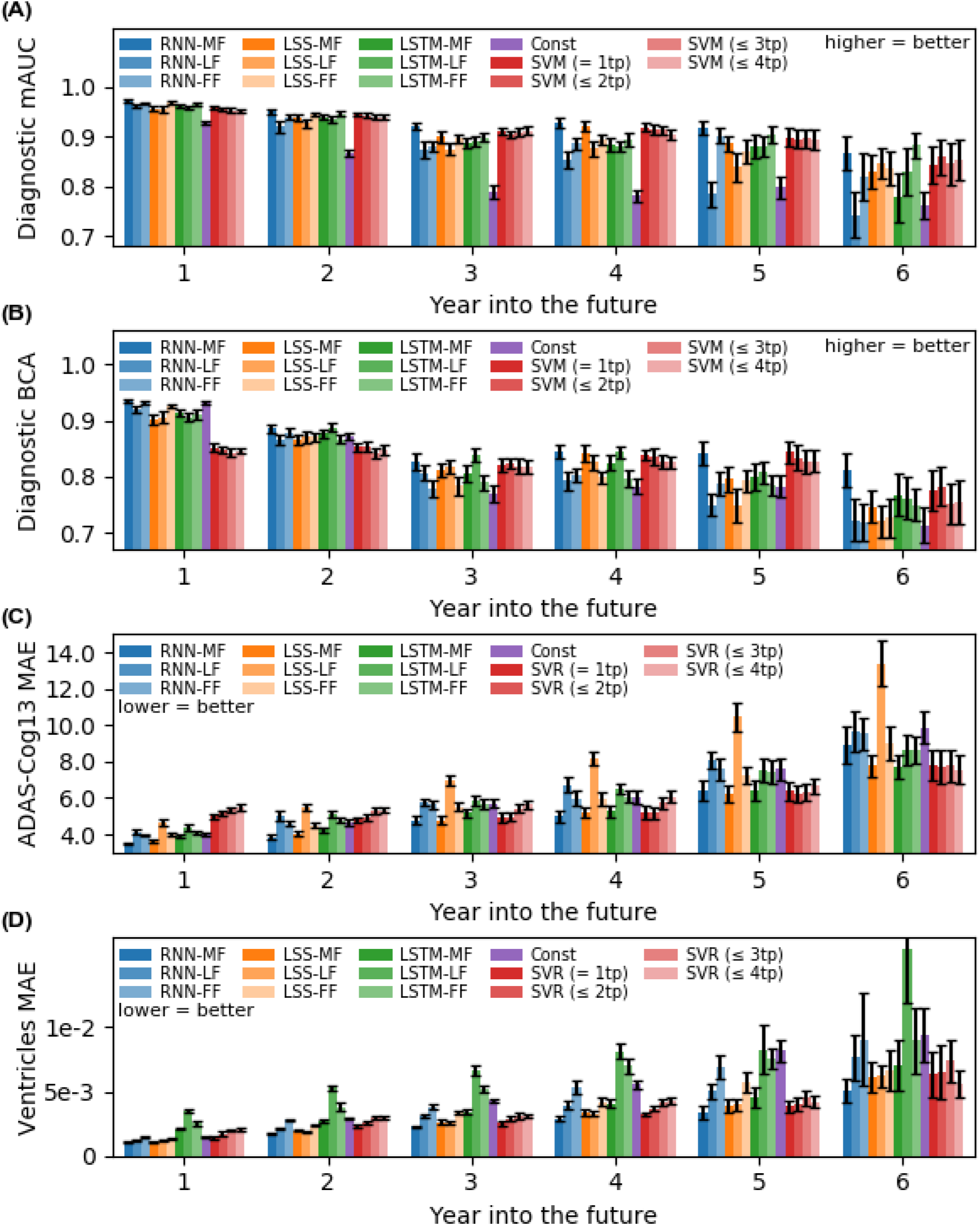
Prediction performance from Figure S1 broken down in yearly interval up to 6 years into the future. For clinical diagnosis, higher mAUC and BCA values indicate better performance. For ADAS-Cog13 and Ventricles, lower MAE indicates better performance. FF indicates forward filling. LF indicates linear filling. MF indicates model filling. SVM/SVR (= 1tp) utilized one input timepoint. SVM/SVR (≤ 2tp) utilized at most 2 input timepoints (see Section 2.5.2 for details) and so on.

**Table S1.** MinimalRNN hyper-parameters (Forward filling)

| Fold No. | Input dropout | Recurrent dropout | Learning rate | Layer size | No. layers | Weight decay |
| --- | --- | --- | --- | --- | --- | --- |
| 1 | 0.5 | 0.5 | 0.006979 | 256 | 3 | $10^{-6}$ |
| 2 | 0.2 | 0.0 | 0.000316 | 256 | 3 | $10^{-7}$ |
| 3 | 0.0 | 0.0 | 0.000118 | 128 | 3 | $10^{-4}$ |
| 4 | 0.2 | 0.0 | 0.000074 | 128 | 3 | $10^{-4}$ |
| 5 | 0.3 | 0.3 | 0.000682 | 256 | 1 | $10^{-6}$ |
| 6 | 0.2 | 0.4 | 0.000469 | 256 | 1 | $10^{-4}$ |
| 7 | 0.4 | 0.0 | 0.000050 | 256 | 3 | $10^{-5}$ |
| 8 | 0.2 | 0.0 | 0.000118 | 256 | 2 | $10^{-4}$ |
| 9 | 0.0 | 0.0 | 0.000432 | 256 | 2 | $10^{-5}$ |
| 10 | 0.0 | 0.0 | 0.000118 | 256 | 2 | $10^{-5}$ |
| 11 | 0.0 | 0.0 | 0.000118 | 512 | 3 | $10^{-4}$ |
| 12 | 0.5 | 0.4 | 0.003746 | 256 | 3 | $10^{-6}$ |
| 13 | 0.1 | 0.1 | 0.000163 | 256 | 2 | $10^{-5}$ |
| 14 | 0.4 | 0.5 | 0.000649 | 256 | 1 | $10^{-7}$ |
| 15 | 0.2 | 0.4 | 0.000351 | 128 | 3 | $10^{-6}$ |
| 16 | 0.5 | 0.3 | 0.006105 | 128 | 3 | $10^{-6}$ |
| 17 | 0.3 | 0.4 | 0.000703 | 512 | 1 | $10^{-7}$ |
| 18 | 0.3 | 0.0 | 0.000118 | 128 | 2 | $10^{-5}$ |
| 19 | 0.1 | 0.2 | 0.000090 | 128 | 3 | $10^{-4}$ |
| 20 | 0.1 | 0.1 | 0.000224 | 256 | 2 | $10^{-5}$ |
| mean±std | **0.23±0.17** | **0.18±0.20** | **$(1.1±2.0) \times 10^{-3}$** | **243±109** | **2.30±0.80** | **$(3.3±4.0) \times 10^{-5}$** |

**Table S2.**
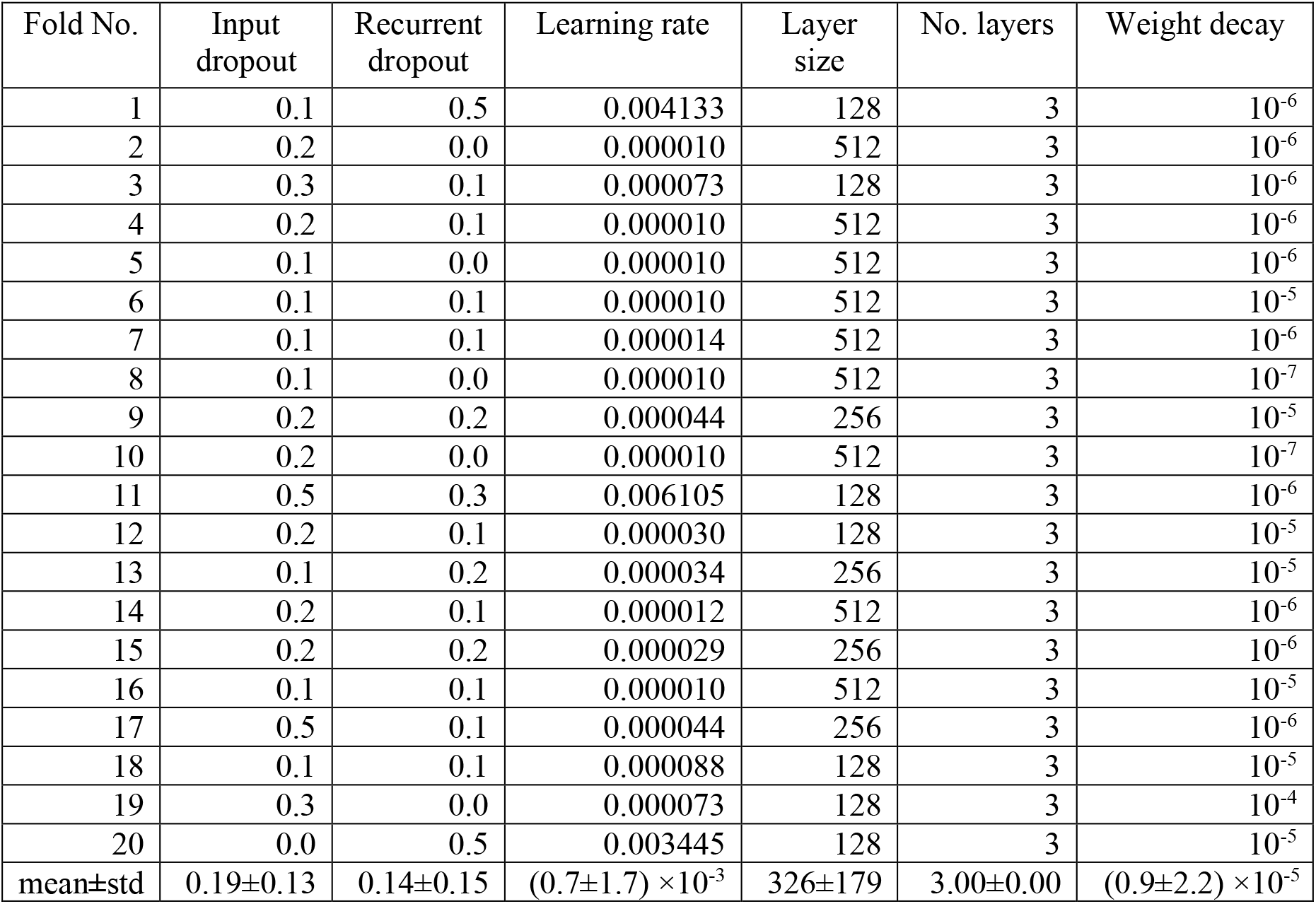
MinimalRNN hyper-parameters (Linear filling)

**Table S3.** MinimalRNN hyper-parameters (Model filling)

| Fold No. | Input dropout | Recurrent dropout | Learning rate | Layer size | No. layers | Weight decay |
| --- | --- | --- | --- | --- | --- | --- |
| 1 | 0.1 | 0.4 | 0.00129 | 128 | 2 | $10^{-5}$ |
| 2 | 0.1 | 0.4 | 0.00133 | 128 | 2 | $10^{-7}$ |
| 3 | 0.4 | 0.4 | 0.00090 | 128 | 1 | $10^{-5}$ |
| 4 | 0.1 | 0.4 | 0.00084 | 128 | 2 | $10^{-6}$ |
| 5 | 0.1 | 0.5 | 0.00084 | 128 | 2 | $10^{-5}$ |
| 6 | 0.4 | 0.3 | 0.00084 | 128 | 1 | $10^{-5}$ |
| 7 | 0.0 | 0.2 | 0.00084 | 128 | 2 | $10^{-4}$ |
| 8 | 0.1 | 0.1 | 0.00094 | 128 | 2 | $10^{-7}$ |
| 9 | 0.4 | 0.4 | 0.00035 | 128 | 2 | $10^{-7}$ |
| 10 | 0.1 | 0.3 | 0.00084 | 128 | 2 | $10^{-6}$ |
| 11 | 0.0 | 0.2 | 0.00084 | 128 | 3 | $10^{-6}$ |
| 12 | 0.0 | 0.2 | 0.00157 | 128 | 2 | $10^{-4}$ |
| 13 | 0.0 | 0.5 | 0.00021 | 128 | 2 | $10^{-6}$ |
| 14 | 0.2 | 0.4 | 0.00147 | 128 | 1 | $10^{-7}$ |
| 15 | 0.1 | 0.3 | 0.00177 | 128 | 2 | $10^{-6}$ |
| 16 | 0.1 | 0.4 | 0.00084 | 128 | 2 | $10^{-7}$ |
| 17 | 0.1 | 0.5 | 0.00084 | 128 | 1 | $10^{-4}$ |
| 18 | 0.0 | 0.4 | 0.00058 | 128 | 2 | $10^{-4}$ |
| 19 | 0.2 | 0.5 | 0.00811 | 128 | 2 | $10^{-7}$ |
| 20 | 0.1 | 0.2 | 0.00130 | 128 | 2 | $10^{-7}$ |
| mean±std | 0.13±0.13 | 0.35±0.12 | $(1.3\pm1.6)\times10^{-3}$ | 128±0 | 1.85±0.49 | $(2.2\pm4.0)\times10^{-7}$ |

**Table S4.** LSS hyper-parameters (Forward filling)

| Fold No. | Input dropout | Recurrent dropout | Learning rate | Layer size | No. layers | Weight decay |
| --- | --- | --- | --- | --- | --- | --- |
| 1 | 0.1 | 0.3 | 0.000207 | 128 | 3 | $10^{-4}$ |
| 2 | 0.0 | 0.0 | 0.000118 | 256 | 2 | $10^{-5}$ |
| 3 | 0.1 | 0.2 | 0.000316 | 256 | 3 | $10^{-7}$ |
| 4 | 0.5 | 0.1 | 0.000838 | 128 | 1 | $10^{-5}$ |
| 5 | 0.2 | 0.3 | 0.000316 | 256 | 1 | $10^{-7}$ |
| 6 | 0.5 | 0.0 | 0.000118 | 128 | 2 | $10^{-5}$ |
| 7 | 0.0 | 0.0 | 0.000118 | 256 | 2 | $10^{-5}$ |
| 8 | 0.2 | 0.1 | 0.000848 | 256 | 3 | $10^{-4}$ |
| 9 | 0.1 | 0.0 | 0.000062 | 512 | 1 | $10^{-7}$ |
| 10 | 0.0 | 0.0 | 0.000039 | 512 | 2 | $10^{-6}$ |
| 11 | 0.4 | 0.2 | 0.000316 | 256 | 1 | $10^{-6}$ |
| 12 | 0.2 | 0.0 | 0.000848 | 128 | 2 | $10^{-4}$ |
| 13 | 0.2 | 0.0 | 0.000480 | 512 | 1 | $10^{-4}$ |
| 14 | 0.2 | 0.2 | 0.000316 | 256 | 1 | $10^{-7}$ |
| 15 | 0.0 | 0.1 | 0.000118 | 256 | 3 | $10^{-7}$ |
| 16 | 0.2 | 0.0 | 0.000072 | 128 | 3 | $10^{-5}$ |
| 17 | 0.3 | 0.0 | 0.000073 | 128 | 2 | $10^{-4}$ |
| 18 | 0.3 | 0.1 | 0.000054 | 256 | 2 | $10^{-5}$ |
| 19 | 0.0 | 0.3 | 0.000501 | 128 | 2 | $10^{-5}$ |
| 20 | 0.0 | 0.0 | 0.000118 | 256 | 2 | $10^{-5}$ |
| mean±std | 0.18±0.16 | 0.10±0.11 | $(3.0\pm2.7)\times10^{-4}$ | 250±128 | 1.95±0.76 | $(2.9\pm4.2)\times10^{-5}$ |

**Table S5.** LSS hyper-parameters (Linear filling)

| Fold No. | Input dropout | Recurrent dropout | Learning rate | Layer size | No. layers | Weight decay |
| --- | --- | --- | --- | --- | --- | --- |
| 1 | 0.3 | 0.0 | 0.000011 | 512 | 3 | $10^{-4}$ |
| 2 | 0.2 | 0.1 | 0.000097 | 128 | 2 | $10^{-5}$ |
| 3 | 0.1 | 0.1 | 0.002450 | 128 | 2 | $10^{-7}$ |
| 4 | 0.0 | 0.2 | 0.002242 | 128 | 3 | $10^{-7}$ |
| 5 | 0.4 | 0.0 | 0.000011 | 512 | 3 | $10^{-5}$ |
| 6 | 0.5 | 0.2 | 0.000078 | 128 | 3 | $10^{-7}$ |
| 7 | 0.2 | 0.0 | 0.000011 | 512 | 3 | $10^{-7}$ |
| 8 | 0.1 | 0.0 | 0.000012 | 512 | 3 | $10^{-5}$ |
| 9 | 0.1 | 0.1 | 0.000316 | 256 | 1 | $10^{-7}$ |
| 10 | 0.4 | 0.0 | 0.000010 | 512 | 3 | $10^{-6}$ |
| 11 | 0.4 | 0.1 | 0.000097 | 128 | 1 | $10^{-7}$ |
| 12 | 0.3 | 0.1 | 0.000316 | 128 | 1 | $10^{-5}$ |
| 13 | 0.2 | 0.0 | 0.000084 | 128 | 1 | $10^{-7}$ |
| 14 | 0.5 | 0.1 | 0.000067 | 128 | 3 | $10^{-7}$ |
| 15 | 0.2 | 0.1 | 0.000065 | 128 | 2 | $10^{-7}$ |
| 16 | 0.4 | 0.1 | 0.000096 | 128 | 1 | $10^{-6}$ |
| 17 | 0.3 | 0.1 | 0.000118 | 128 | 2 | $10^{-5}$ |
| 18 | 0.2 | 0.1 | 0.004458 | 128 | 2 | $10^{-7}$ |
| 19 | 0.2 | 0.0 | 0.000011 | 256 | 2 | $10^{-7}$ |
| 20 | 0.2 | 0.1 | 0.000316 | 128 | 1 | $10^{-6}$ |
| mean±std | 0.26±0.14 | 0.08±0.06 | $(0.5\pm1.2)\times10^{-3}$ | 237±168 | 2.10±0.85 | $(0.8\pm2.2)\times10^{-5}$ |

**Table S6.** LSS hyper-parameters (Model filling)

| Fold No. | Input dropout | Recurrent dropout | Learning rate | Layer size | No. layers | Weight decay |
| --- | --- | --- | --- | --- | --- | --- |
| 1 | 0.4 | 0.1 | 0.00072 | 128 | 2 | $10^{-5}$ |
| 2 | 0.0 | 0.2 | 0.00056 | 128 | 3 | $10^{-6}$ |
| 3 | 0.2 | 0.0 | 0.00031 | 512 | 1 | $10^{-7}$ |
| 4 | 0.2 | 0.1 | 0.00084 | 128 | 2 | $10^{-6}$ |
| 5 | 0.2 | 0.0 | 0.00031 | 512 | 1 | $10^{-7}$ |
| 6 | 0.1 | 0.1 | 0.00041 | 256 | 1 | $10^{-7}$ |
| 7 | 0.1 | 0.0 | 0.00022 | 512 | 1 | $10^{-5}$ |
| 8 | 0.2 | 0.0 | 0.00031 | 256 | 3 | $10^{-7}$ |
| 9 | 0.0 | 0.0 | 0.00010 | 256 | 3 | $10^{-6}$ |
| 10 | 0.2 | 0.2 | 0.00084 | 128 | 2 | $10^{-6}$ |
| 11 | 0.0 | 0.0 | 0.00011 | 512 | 2 | $10^{-4}$ |
| 12 | 0.2 | 0.2 | 0.00031 | 256 | 1 | $10^{-7}$ |
| 13 | 0.1 | 0.2 | 0.00031 | 256 | 1 | $10^{-5}$ |
| 14 | 0.5 | 0.2 | 0.00066 | 128 | 3 | $10^{-7}$ |
| 15 | 0.0 | 0.0 | 0.00019 | 256 | 3 | $10^{-5}$ |
| 16 | 0.0 | 0.2 | 0.00009 | 256 | 3 | $10^{-6}$ |
| 17 | 0.0 | 0.2 | 0.00031 | 256 | 1 | $10^{-5}$ |
| 18 | 0.0 | 0.0 | 0.00011 | 256 | 2 | $10^{-5}$ |
| 19 | 0.2 | 0.5 | 0.00084 | 128 | 2 | $10^{-6}$ |
| 20 | 0.3 | 0.5 | 0.00084 | 128 | 3 | $10^{-4}$ |
| mean±std | 0.15±0.14 | 0.14±0.15 | $(4.3\pm2.8)\times10^{-4}$ | 262±140 | 2.00±0.85 | $(1.3\pm3.0)\times10^{-5}$ |

**Table S7.** LSTM hyper-parameters (Forward filling)

| Fold No. | Input dropout | Recurrent dropout | Learning rate | Layer size | No. layers | Weight decay |
| --- | --- | --- | --- | --- | --- | --- |
| 1 | 0.0 | 0.0 | 0.000699 | 256 | 1 | $10^{-7}$ |
| 2 | 0.0 | 0.0 | 0.000035 | 256 | 3 | $10^{-5}$ |
| 3 | 0.1 | 0.0 | 0.000740 | 128 | 1 | $10^{-5}$ |
| 4 | 0.2 | 0.1 | 0.000058 | 512 | 2 | $10^{-6}$ |
| 5 | 0.0 | 0.0 | 0.000721 | 512 | 1 | $10^{-7}$ |
| 6 | 0.0 | 0.0 | 0.000074 | 512 | 2 | $10^{-5}$ |
| 7 | 0.1 | 0.2 | 0.001379 | 128 | 2 | $10^{-7}$ |
| 8 | 0.0 | 0.0 | 0.000175 | 512 | 1 | $10^{-7}$ |
| 9 | 0.0 | 0.0 | 0.000320 | 512 | 1 | $10^{-6}$ |
| 10 | 0.0 | 0.0 | 0.000059 | 512 | 1 | $10^{-7}$ |
| 11 | 0.1 | 0.1 | 0.000096 | 512 | 1 | $10^{-4}$ |
| 12 | 0.2 | 0.2 | 0.000117 | 256 | 1 | $10^{-6}$ |
| 13 | 0.1 | 0.0 | 0.000200 | 512 | 2 | $10^{-6}$ |
| 14 | 0.0 | 0.0 | 0.000373 | 512 | 2 | $10^{-7}$ |
| 15 | 0.1 | 0.0 | 0.000059 | 512 | 1 | $10^{-4}$ |
| 16 | 0.0 | 0.0 | 0.000080 | 256 | 2 | $10^{-5}$ |
| 17 | 0.1 | 0.0 | 0.000643 | 256 | 1 | $10^{-6}$ |
| 18 | 0.0 | 0.0 | 0.000451 | 512 | 1 | $10^{-4}$ |
| 19 | 0.0 | 0.0 | 0.000090 | 512 | 1 | $10^{-4}$ |
| 20 | 0.1 | 0.0 | 0.000432 | 512 | 1 | $10^{-6}$ |
| mean±std | 0.06±0.07 | 0.03±0.07 | $(3.4\pm3.5)\times10^{-4}$ | 410±147 | 1.40±0.60 | $(2.7\pm4.3)\times10^{-5}$ |

**Table S8.** LSTM hyper-parameters (Linear filling)

| Fold No. | Input dropout | Recurrent dropout | Learning rate | Layer size | No. layers | Weight decay |
| --- | --- | --- | --- | --- | --- | --- |
| 1 | 0.2 | 0.2 | 0.000208 | 256 | 1 | $10^{-7}$ |
| 2 | 0.2 | 0.2 | 0.000541 | 128 | 1 | $10^{-7}$ |
| 3 | 0.2 | 0.2 | 0.000237 | 256 | 1 | $10^{-6}$ |
| 4 | 0.2 | 0.2 | 0.000058 | 256 | 3 | $10^{-7}$ |
| 5 | 0.2 | 0.2 | 0.000316 | 256 | 1 | $10^{-7}$ |
| 6 | 0.2 | 0.2 | 0.000303 | 256 | 1 | $10^{-7}$ |
| 7 | 0.2 | 0.2 | 0.000130 | 512 | 1 | $10^{-6}$ |
| 8 | 0.1 | 0.1 | 0.000021 | 512 | 3 | $10^{-6}$ |
| 9 | 0.2 | 0.2 | 0.000650 | 256 | 1 | $10^{-6}$ |
| 10 | 0.1 | 0.2 | 0.000074 | 256 | 1 | $10^{-7}$ |
| 11 | 0.1 | 0.1 | 0.000080 | 256 | 1 | $10^{-6}$ |
| 12 | 0.2 | 0.2 | 0.000161 | 256 | 1 | $10^{-7}$ |
| 13 | 0.2 | 0.2 | 0.000237 | 256 | 1 | $10^{-6}$ |
| 14 | 0.2 | 0.2 | 0.000168 | 256 | 1 | $10^{-7}$ |
| 15 | 0.2 | 0.2 | 0.000101 | 128 | 3 | $10^{-7}$ |
| 16 | 0.2 | 0.2 | 0.000072 | 256 | 2 | $10^{-7}$ |
| 17 | 0.2 | 0.2 | 0.000119 | 256 | 1 | $10^{-6}$ |
| 18 | 0.2 | 0.2 | 0.000190 | 256 | 1 | $10^{-7}$ |
| 19 | 0.2 | 0.2 | 0.000191 | 256 | 1 | $10^{-7}$ |
| 20 | 0.2 | 0.2 | 0.000149 | 256 | 1 | $10^{-6}$ |
| mean±std | 0.19±0.04 | 0.19±0.03 | $(2.0\pm1.6)\times10^{-4}$ | 269±92 | 1.35±7.45 | $(4.6\pm4.5)\times10^{-7}$ |

**Table S9.** LSTM hyper-parameters (Model filling)

| Fold No. | Input dropout | Recurrent dropout | Learning rate | Layer size | No. layers | Weight decay |
| --- | --- | --- | --- | --- | --- | --- |
| 1 | 0.1 | 0.1 | 0.002276 | 512 | 1 | $10^{-4}$ |
| 2 | 0.0 | 0.1 | 0.010000 | 128 | 1 | $10^{-7}$ |
| 3 | 0.1 | 0.0 | 0.006868 | 256 | 1 | $10^{-4}$ |
| 4 | 0.1 | 0.3 | 0.010000 | 512 | 1 | $10^{-5}$ |
| 5 | 0.1 | 0.4 | 0.008537 | 256 | 1 | $10^{-4}$ |
| 6 | 0.2 | 0.0 | 0.002276 | 256 | 1 | $10^{-5}$ |
| 7 | 0.4 | 0.1 | 0.005509 | 256 | 2 | $10^{-4}$ |
| 8 | 0.1 | 0.4 | 0.010000 | 128 | 2 | $10^{-4}$ |
| 9 | 0.1 | 0.1 | 0.005038 | 128 | 1 | $10^{-7}$ |
| 10 | 0.2 | 0.1 | 0.003823 | 128 | 1 | $10^{-5}$ |
| 11 | 0.1 | 0.4 | 0.010000 | 256 | 1 | $10^{-7}$ |
| 12 | 0.2 | 0.1 | 0.009419 | 128 | 1 | $10^{-4}$ |
| 13 | 0.0 | 0.4 | 0.010000 | 128 | 1 | $10^{-7}$ |
| 14 | 0.2 | 0.1 | 0.010000 | 128 | 1 | $10^{-4}$ |
| 15 | 0.2 | 0.0 | 0.003699 | 512 | 1 | $10^{-7}$ |
| 16 | 0.0 | 0.3 | 0.005813 | 128 | 1 | $10^{-7}$ |
| 17 | 0.2 | 0.0 | 0.004785 | 128 | 1 | $10^{-5}$ |
| 18 | 0.2 | 0.0 | 0.010000 | 128 | 1 | $10^{-7}$ |
| 19 | 0.1 | 0.2 | 0.001827 | 128 | 1 | $10^{-6}$ |
| 20 | 0.1 | 0.1 | 0.006821 | 128 | 1 | $10^{-4}$ |
| mean±std | 0.14±0.09 | 0.16±0.15 | $(6.8\pm3.0)\times10^{-3}$ | 218±138 | 1.10±0.31 | $(4.2\pm4.9)\times10^{-5}$ |

**Table S10a.** SVR hyper-parameters (1 input timepoint). Recall that we trained separate SVM/SVR models to predict 10 sets of timepoints (spaced 6 months apart) into the future, i.e., 6, 12, 18, …, 60 months into the future. We note that the hyperparameters are the same across these 10 sets of SVM/SVR models. The reason is to avoid an explosion in the number of hyperparameters.

|  | ADAS-Cog13 |  |  |  | Ventricles |  |  |  |
| --- | --- | --- | --- | --- | --- | --- | --- | --- |
| Fold No. | C | Epsilon | Gamma | Kernel | C | Epsilon | Gamma | Kernel |
| 1 | 1.4E-1 | 1.2E-3 | 1.6E-2 | rbf | 1.4E-0 | 1.4E-2 | 2.0E-1 | linear |
| 2 | 3.9E-2 | 8.7E-2 | 1.8E-3 | linear | 8.7E-1 | 7.6E-2 | 1.0E-3 | linear |
| 3 | 1.1E-0 | 2.6E-2 | 1.5E-2 | rbf | 8.4E-1 | 5.7E-2 | 5.1E-1 | linear |
| 4 | 2.3E-0 | 1.1E-3 | 3.4E-3 | rbf | 2.0E-0 | 8.7E-3 | 1.8E-0 | linear |
| 5 | 4.8E-1 | 2.2E-3 | 4.7E-1 | linear | 2.5E-0 | 8.4E-3 | 1.3E-3 | rbf |
| 6 | 4.9E-1 | 6.7E-3 | 1.2E-0 | linear | 7.8E-1 | 1.3E-2 | 7.7E-2 | linear |
| 7 | 1.3E-1 | 2.9E-1 | 2.4E-1 | linear | 1.2E-1 | 1.0E-3 | 2.1E-3 | linear |
| 8 | 1.6E-3 | 1.0E-3 | 4.5E-3 | linear | 2.6E-0 | 2.2E-2 | 1.6E-3 | linear |
| 9 | 7.9E-1 | 2.2E-2 | 6.5E-3 | rbf | 7.8E-1 | 4.9E-3 | 3.4E-1 | linear |
| 10 | 2.9E-1 | 2.6E-3 | 1.4E-2 | rbf | 2.8E-1 | 1.0E-3 | 5.8E-1 | linear |
| 11 | 3.7E-0 | 3.3E-3 | 4.6E-3 | rbf | 6.4E-1 | 3.7E-2 | 4.0E-0 | linear |
| 12 | 3.1E-1 | 1.2E-1 | 4.6E-3 | rbf | 6.7E-1 | 6.5E-2 | 1.2E-2 | linear |
| 13 | 2.3E-2 | 1.6E-2 | 5.9E-3 | linear | 9.3E-0 | 1.6E-2 | 6.9E-3 | rbf |
| 14 | 5.8E-0 | 1.0E-2 | 5.4E-3 | rbf | 1.3E-0 | 3.3E-2 | 5.2E+1 | linear |
| 15 | 3.2E-1 | 9.7E-3 | 1.0E-3 | linear | 2.8E-2 | 3.5E-2 | 3.9E+2 | linear |
| 16 | 4.4E-1 | 5.0E-1 | 1.5E-3 | linear | 1.6E-2 | 1.0E-3 | 1.7E+1 | linear |
| 17 | 4.1E-2 | 1.6E-1 | 2.7E+2 | linear | 3.8E+1 | 1.3E-3 | 2.3E-3 | rbf |
| 18 | 2.4E-1 | 1.2E-2 | 6.1E+1 | linear | 1.3E-2 | 2.3E-2 | 2.6E-3 | linear |
| 19 | 7.8E-3 | 1.0E-2 | 7.6E-1 | linear | 7.4E-2 | 1.0E-3 | 2.3E-3 | linear |
| 20 | 4.8E-1 | 1.0E-3 | 1.0E-3 | linear | 3.0E-1 | 2.1E-3 | 3.7E-1 | linear |
| mean | 8.6E-1 | 6.4E-2 | 1.7E+1 |  | 3.1E-0 | 2.1E-2 | 2.4E+1 |  |
| std | 1.5E-0 | 1.3E-1 | 6.1E+1 |  | 8.4E-0 | 2.3E-2 | 8.8E+1 |  |

**Table S10b.** SVM hyper-parameters (1 input timepoint). Recall that we trained separate SVM/SVR models to predict 10 sets of timepoints (spaced 6 months apart) into the future, i.e., 6, 12, 18, …, 60 months into the future. We note that the hyperparameters are the same across these 10 sets of SVM/SVR models. The reason is to avoid an explosion in the number of hyperparameters.

|  | Diagnosis |  |  |  |  |  |  |  |
| --- | --- | --- | --- | --- | --- | --- | --- | --- |
| Fold No. | C | Gamma | Kernel |  | Fold No. | C | Gamma | Kernel |
| 1 | 2.9E-2 | 1.0E-2 | linear |  | 11 | 4.2E-3 | 1.3E-2 | linear |
| 2 | 3.9E-3 | 3.6E-3 | linear |  | 12 | 1.2E-1 | 1.2E-0 | linear |
| 3 | 2.6E-3 | 8.5E-3 | linear |  | 13 | 6.6E-3 | 2.0E-1 | linear |
| 4 | 6.3E-3 | 2.1E-3 | linear |  | 14 | 4.2E-3 | 2.5E-2 | linear |
| 5 | 1.5E-2 | 2.4E-3 | linear |  | 15 | 5.5E-3 | 1.2E-3 | linear |
| 6 | 5.9E-3 | 2.2E-3 | linear |  | 16 | 2.1E-2 | 7.2E+1 | linear |
| 7 | 3.3E-3 | 1.1E-2 | linear |  | 17 | 8.1E-2 | 2.6E-2 | linear |
| 8 | 4.4E-1 | 4.8E-3 | linear |  | 18 | 8.9E-1 | 2.8E-3 | linear |
| 9 | 7.3E-1 | 1.7E-2 | rbf |  | 19 | 8.5E-2 | 2.9E-2 | linear |
| 10 | 7.4E-3 | 1.2E-3 | linear |  | 20 | 6.0E-3 | 7.8E-0 | linear |
|  |  |  |  |  | mean | 1.2E-01 | 4.1E+0 |  |
|  |  |  |  |  | Std | 2.6E-01 | 1.6E+1 |  |

**Table S11a.**
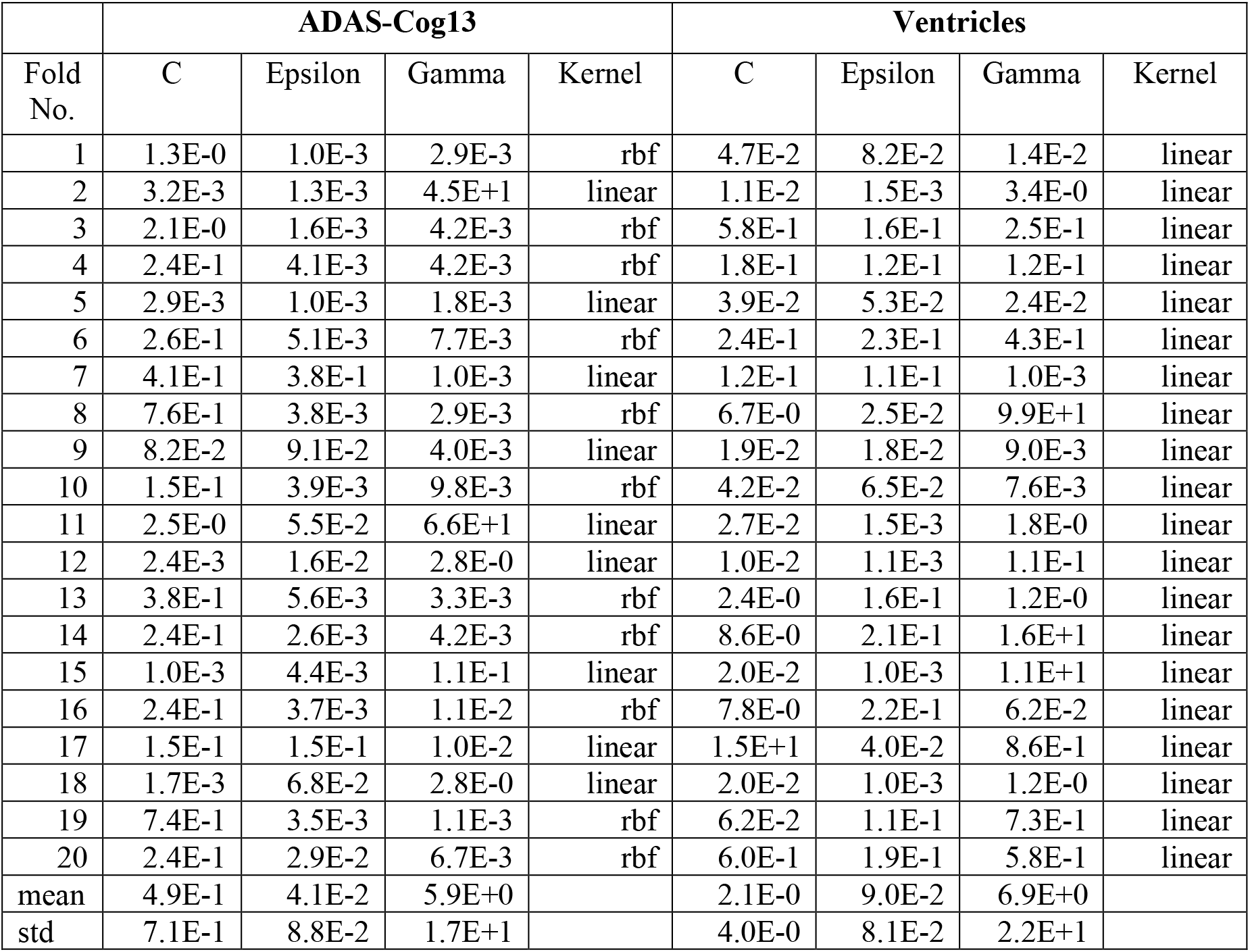
SVR hyper-parameters (2 input timepoints). Recall that we trained separate SVM/SVR models to predict 10 sets of timepoints (spaced 6 months apart) into the future, i.e., 6, 12, 18, …, 60 months into the future. We note that the hyperparameters are the same across these 10 sets of SVM/SVR models. The reason is to avoid an explosion in the number of hyperparameters.

|  | ADAS-Cog13 |  |  |  | Ventricles |  |  |  |
| --- | --- | --- | --- | --- | --- | --- | --- | --- |
| Fold No. | C | Epsilon | Gamma | Kernel | C | Epsilon | Gamma | Kernel |
| 1 | 1.3E-0 | 1.0E-3 | 2.9E-3 | rbf | 4.7E-2 | 8.2E-2 | 1.4E-2 | linear |
| 2 | 3.2E-3 | 1.3E-3 | 4.5E+1 | linear | 1.1E-2 | 1.5E-3 | 3.4E-0 | linear |
| 3 | 2.1E-0 | 1.6E-3 | 4.2E-3 | rbf | 5.8E-1 | 1.6E-1 | 2.5E-1 | linear |
| 4 | 2.4E-1 | 4.1E-3 | 4.2E-3 | rbf | 1.8E-1 | 1.2E-1 | 1.2E-1 | linear |
| 5 | 2.9E-3 | 1.0E-3 | 1.8E-3 | linear | 3.9E-2 | 5.3E-2 | 2.4E-2 | linear |
| 6 | 2.6E-1 | 5.1E-3 | 7.7E-3 | rbf | 2.4E-1 | 2.3E-1 | 4.3E-1 | linear |
| 7 | 4.1E-1 | 3.8E-1 | 1.0E-3 | linear | 1.2E-1 | 1.1E-1 | 1.0E-3 | linear |
| 8 | 7.6E-1 | 3.8E-3 | 2.9E-3 | rbf | 6.7E-0 | 2.5E-2 | 9.9E+1 | linear |
| 9 | 8.2E-2 | 9.1E-2 | 4.0E-3 | linear | 1.9E-2 | 1.8E-2 | 9.0E-3 | linear |
| 10 | 1.5E-1 | 3.9E-3 | 9.8E-3 | rbf | 4.2E-2 | 6.5E-2 | 7.6E-3 | linear |
| 11 | 2.5E-0 | 5.5E-2 | 6.6E+1 | linear | 2.7E-2 | 1.5E-3 | 1.8E-0 | linear |
| 12 | 2.4E-3 | 1.6E-2 | 2.8E-0 | linear | 1.0E-2 | 1.1E-3 | 1.1E-1 | linear |
| 13 | 3.8E-1 | 5.6E-3 | 3.3E-3 | rbf | 2.4E-0 | 1.6E-1 | 1.2E-0 | linear |
| 14 | 2.4E-1 | 2.6E-3 | 4.2E-3 | rbf | 8.6E-0 | 2.1E-1 | 1.6E+1 | linear |
| 15 | 1.0E-3 | 4.4E-3 | 1.1E-1 | linear | 2.0E-2 | 1.0E-3 | 1.1E+1 | linear |
| 16 | 2.4E-1 | 3.7E-3 | 1.1E-2 | rbf | 7.8E-0 | 2.2E-1 | 6.2E-2 | linear |
| 17 | 1.5E-1 | 1.5E-1 | 1.0E-2 | linear | 1.5E+1 | 4.0E-2 | 8.6E-1 | linear |
| 18 | 1.7E-3 | 6.8E-2 | 2.8E-0 | linear | 2.0E-2 | 1.0E-3 | 1.2E-0 | linear |
| 19 | 7.4E-1 | 3.5E-3 | 1.1E-3 | rbf | 6.2E-2 | 1.1E-1 | 7.3E-1 | linear |
| 20 | 2.4E-1 | 2.9E-2 | 6.7E-3 | rbf | 6.0E-1 | 1.9E-1 | 5.8E-1 | linear |
| mean | 4.9E-1 | 4.1E-2 | 5.9E+0 |  | 2.1E-0 | 9.0E-2 | 6.9E+0 |  |
| std | 7.1E-1 | 8.8E-2 | 1.7E+1 |  | 4.0E-0 | 8.1E-2 | 2.2E+1 |  |

**Table S11b.** SVM hyper-parameters (2 input timepoints). Recall that we trained separate SVM/SVR models to predict 10 sets of timepoints (spaced 6 months apart) into the future, i.e., 6, 12, 18, …, 60 months into the future. We note that the hyperparameters are the same across these 10 sets of SVM/SVR models. The reason is to avoid an explosion in the number of hyperparameters.

|  | Diagnosis |  |  |  |  |  |  |  |
| --- | --- | --- | --- | --- | --- | --- | --- | --- |
| Fold No. | C | Gamma | Kernel |  | Fold No. | C | Gamma | Kernel |
| 1 | 4.5E-1 | 3.3E+2 | linear |  | 11 | 5.4E-3 | 1.5E-3 | linear |
| 2 | 1.0E-3 | 2.3E-2 | linear |  | 12 | 2.4E-1 | 6.8E-3 | linear |
| 3 | 4.6E-3 | 2.8E-1 | linear |  | 13 | 9.6E-3 | 6.3E-1 | linear |
| 4 | 2.4E-3 | 1.3E-1 | linear |  | 14 | 2.4E-3 | 6.7E-3 | linear |
| 5 | 1.8E-2 | 1.1E-1 | linear |  | 15 | 2.2E-2 | 2.2E+2 | linear |
| 6 | 1.4E-1 | 3.6E-2 | linear |  | 16 | 1.1E-1 | 2.8E-3 | linear |
| 7 | 1.9E-2 | 3.9E-2 | linear |  | 17 | 5.4E-2 | 1.1E+2 | linear |
| 8 | 2.8E-2 | 7.1E-2 | linear |  | 18 | 1.9E-1 | 6.5E-0 | linear |
| 9 | 3.6E-2 | 5.5E+1 | linear |  | 19 | 8.7E-2 | 5.1E-3 | linear |
| 10 | 3.5E-3 | 2.0E-1 | linear |  | 20 | 4.4E-3 | 7.2E-2 | linear |
|  |  |  |  |  | mean | 7.2E-2 | 3.6E+1 |  |
|  |  |  |  |  | std | 1.1E-1 | 8.8E+1 |  |

**Table 12a.** SVR hyper-parameters (3 input timepoints). Recall that we trained separate SVM/SVR models to predict 10 sets of timepoints (spaced 6 months apart) into the future, i.e., 6, 12, 18, …, 60 months into the future. We note that the hyperparameters are the same across these 10 sets of SVM/SVR models. The reason is to avoid an explosion in the number of hyperparameters.

|  | ADAS-Cog13 |  |  |  | Ventricles |  |  |  |
| --- | --- | --- | --- | --- | --- | --- | --- | --- |
| Fold No. | C | Epsilon | Gamma | Kernel | C | Epsilon | Gamma | Kernel |
| 1 | 2.3E-2 | 5.4E-1 | 2.2E-3 | linear | 1.2E-2 | 2.5E-3 | 1.1E-2 | linear |
| 2 | 3.3E-3 | 1.1E-1 | 1.0E-3 | linear | 1.1E-2 | 1.0E-3 | 1.4E-0 | linear |
| 3 | 2.9E-3 | 1.6E-3 | 3.8E+1 | linear | 1.7E-2 | 3.3E-2 | 1.8E-2 | linear |
| 4 | 8.0E-0 | 2.5E-1 | 1.0E-3 | rbf | 4.9E-2 | 9.1E-2 | 7.4E-3 | linear |
| 5 | 1.5E-3 | 1.0E-3 | 5.0E+1 | linear | 6.3E+1 | 8.9E-2 | 1.0E-3 | rbf |
| 6 | 6.5E-2 | 4.2E-3 | 9.4E-3 | rbf | 3.5E-2 | 8.1E-2 | 1.8E-2 | linear |
| 7 | 9.1E-2 | 5.7E-1 | 1.1E-2 | linear | 4.0E-2 | 3.4E-2 | 1.4E+1 | linear |
| 8 | 4.5E-1 | 1.0E-2 | 4.2E-3 | rbf | 5.0E-0 | 3.2E-3 | 2.2E-3 | rbf |
| 9 | 2.1E-0 | 6.8E-3 | 1.2E-3 | rbf | 5.6E-1 | 2.2E-1 | 1.1E+1 | linear |
| 10 | 3.0E-1 | 2.6E-3 | 3.6E-3 | rbf | 1.5E-2 | 1.9E-2 | 1.5E-3 | linear |
| 11 | 3.4E-2 | 1.5E-1 | 1.4E-0 | linear | 4.1E-2 | 1.3E-3 | 3.9E-2 | linear |
| 12 | 2.6E-3 | 6.4E-3 | 1.9E-3 | linear | 1.2E+1 | 5.0E-2 | 1.0E-3 | rbf |
| 13 | 1.6E-1 | 4.1E-1 | 1.1E-2 | linear | 1.0E-2 | 1.0E-3 | 1.2E-2 | linear |
| 14 | 5.1E-2 | 6.3E-2 | 1.7E-2 | linear | 1.1E-1 | 9.1E-2 | 3.0E-1 | linear |
| 15 | 2.5E-2 | 2.9E-1 | 1.5E-1 | linear | 3.6E-2 | 6.7E-2 | 3.4E-3 | linear |
| 16 | 2.3E-1 | 5.6E-3 | 8.3E-3 | rbf | 1.8E+1 | 3.0E-2 | 4.2E+1 | linear |
| 17 | 3.2E-2 | 8.5E-2 | 1.4E-2 | linear | 7.1E-2 | 9.6E-2 | 3.0E-3 | linear |
| 18 | 3.2E-1 | 4.3E-3 | 3.0E-3 | rbf | 2.9E-2 | 6.0E-2 | 3.3E-3 | linear |
| 19 | 1.8E-1 | 5.1E-3 | 4.9E-3 | rbf | 2.8E-1 | 2.1E-1 | 7.1E-2 | linear |
| 20 | 2.4E-1 | 3.7E-3 | 4.2E-3 | rbf | 2.4E+1 | 1.4E-1 | 7.5E-2 | linear |
| mean | 6.1E-1 | 1.3E-1 | 4.5E+0 |  | 6.2E+0 | 6.6E-2 | 3.5E-0 |  |
| std | 1.8E-0 | 1.9E-1 | 1.4E+1 |  | 1.5E+1 | 6.5E-2 | 9.9E-0 |  |

**Table 12b.** SVM hyper-parameters (3 input timepoints). Recall that we trained separate SVM/SVR models to predict 10 sets of timepoints (spaced 6 months apart) into the future, i.e., 6, 12, 18, …, 60 months into the future. We note that the hyperparameters are the same across these 10 sets of SVM/SVR models. The reason is to avoid an explosion in the number of hyperparameters.

|  | Diagnosis |  |  |  |  |  |  |  |
| --- | --- | --- | --- | --- | --- | --- | --- | --- |
| Fold No. | C | Gamma | Kernel |  | Fold No. | C | Gamma | Kernel |
| 1 | 1.1E-1 | 1.3E+0 | linear |  | 11 | 4.5E-2 | 1.0E+1 | linear |
| 2 | 4.2E-3 | 5.3E+2 | linear |  | 12 | 5.2E-2 | 4.4E-1 | linear |
| 3 | 4.6E-2 | 5.4E+0 | linear |  | 13 | 2.9E-2 | 2.8E+1 | linear |
| 4 | 5.2E-3 | 7.0E+1 | linear |  | 14 | 2.0E-2 | 1.3E-3 | linear |
| 5 | 3.0E-2 | 3.6E+2 | linear |  | 15 | 7.7E-2 | 1.3E-1 | linear |
| 6 | 4.5E-2 | 1.2E-2 | linear |  | 16 | 3.3E-2 | 4.5E-1 | linear |
| 7 | 2.1E-3 | 1.0E-3 | linear |  | 17 | 6.3E-2 | 6.1E-0 | linear |
| 8 | 4.1E-2 | 2.7E-2 | linear |  | 18 | 1.3E-2 | 6.8E-2 | linear |
| 9 | 2.3E-2 | 3.5E-2 | linear |  | 19 | 8.1E-3 | 9.3E-0 | linear |
| 10 | 5.9E-2 | 5.8E-1 | linear |  | 20 | 8.4E-3 | 1.2E-2 | linear |
|  |  |  |  |  | mean | 3.6E-2 | 5.1E+1 |  |
|  |  |  |  |  | std | 2.7E-2 | 1.4E+2 |  |

**Table 13a.** SVR hyper-parameters (4 input timepoints). Recall that we trained separate SVM/SVR models to predict 10 sets of timepoints (spaced 6 months apart) into the future, i.e., 6, 12, 18, …, 60 months into the future. We note that the hyperparameters are the same across these 10 sets of SVM/SVR models. The reason is to avoid an explosion in the number of hyperparameters.

|  | ADAS-Cog13 |  |  |  | Ventricles |  |  |  |
| --- | --- | --- | --- | --- | --- | --- | --- | --- |
| Fold No. | C | Epsilon | Gamma | Kernel | C | Epsilon | Gamma | Kernel |
| 1 | 6.1E-2 | 6.3E-1 | 8.1E-3 | linear | 3.9E+1 | 2.5E-2 | 1.0E-3 | rbf |
| 2 | 2.0E-2 | 1.7E-1 | 6.9E-2 | linear | 1.7E-2 | 1.5E-2 | 4.0E-3 | linear |
| 3 | 1.3E-2 | 8.7E-2 | 7.3E-2 | linear | 2.0E-2 | 1.9E-2 | 1.0E-1 | linear |
| 4 | 2.5E-3 | 1.6E-3 | 9.2E-3 | linear | 2.2E-2 | 2.8E-2 | 3.3E-3 | linear |
| 5 | 3.6E-3 | 1.6E-1 | 4.6E-3 | linear | 2.2E-2 | 1.0E-3 | 8.5E-2 | linear |
| 6 | 1.7E-1 | 1.6E-3 | 6.7E-3 | rbf | 2.2E-2 | 4.0E-2 | 2.7E-3 | linear |
| 7 | 3.6E-2 | 6.8E-1 | 5.8E-3 | linear | 2.4E-2 | 1.0E-3 | 1.0E-3 | linear |
| 8 | 4.1E-1 | 1.2E-3 | 2.7E-3 | rbf | 4.5E-2 | 1.5E-2 | 4.5E+0 | linear |
| 9 | 2.2E-2 | 5.2E-1 | 2.3E-2 | linear | 2.9E-2 | 2.3E-2 | 7.3E-1 | linear |
| 10 | 6.0E-3 | 2.0E-3 | 1.0E-3 | linear | 2.4E-2 | 5.8E-2 | 2.8E-1 | linear |
| 11 | 1.5E-2 | 1.2E-2 | 2.2E+1 | linear | 4.6E-2 | 2.5E-2 | 1.0E-3 | linear |
| 12 | 1.3E-2 | 3.7E-1 | 2.8E-1 | linear | 5.7E-2 | 1.0E-1 | 1.5E+2 | linear |
| 13 | 2.2E-2 | 4.3E-1 | 2.6E-2 | linear | 1.9E-2 | 1.3E-3 | 1.3E-2 | linear |
| 14 | 4.0E-2 | 5.3E-1 | 2.3E-0 | linear | 1.4E-1 | 1.2E-1 | 2.1E+1 | linear |
| 15 | 1.1E-2 | 9.6E-3 | 6.7E-1 | linear | 1.6E-2 | 1.5E-3 | 1.8E-1 | linear |
| 16 | 1.7E-1 | 1.0E-3 | 5.2E-3 | rbf | 4.0E-2 | 5.6E-2 | 7.3E-1 | linear |
| 17 | 2.9E-2 | 4.7E-1 | 1.6E-2 | linear | 1.2E-2 | 1.0E-3 | 8.0E-3 | linear |
| 18 | 4.1E-3 | 1.9E-2 | 1.4E-3 | linear | 1.7E-2 | 1.0E-3 | 4.4E-2 | linear |
| 19 | 2.2E-1 | 3.7E-3 | 4.2E-3 | rbf | 3.5E+1 | 4.3E-2 | 8.3E-1 | linear |
| 20 | 3.2E-1 | 2.9E-3 | 3.3E-3 | rbf | 6.0E-2 | 7.3E-2 | 6.7E-1 | linear |
| mean | 8.0E-2 | 2.1E-1 | 1.3E-0 |  | 3.7E+0 | 3.2E-2 | 8.8E+0 |  |
| std | 1.2E-1 | 2.5E-1 | 4.8E-0 |  | 1.1E+1 | 3.4E-2 | 3.3E+1 |  |

**Table 13b.** SVM hyper-parameters (4 input timepoints). Recall that we trained separate SVM/SVR models to predict 10 sets of timepoints (spaced 6 months apart) into the future, i.e., 6, 12, 18, …, 60 months into the future. We note that the hyperparameters are the same across these 10 sets of SVM/SVR models. The reason is to avoid an explosion in the number of hyperparameters.

|  | Diagnosis |  |  |  |  |  |  |  |
| --- | --- | --- | --- | --- | --- | --- | --- | --- |
| Fold No. | C | Gamma | Kernel |  | Fold No. | C | Gamma | Kernel |
| 1 | 9.2E-2 | 1.2E+0 | linear |  | 11 | 3.2E-2 | 5.2E-1 | linear |
| 2 | 8.1E-2 | 1.2E+1 | linear |  | 12 | 4.9E-2 | 1.0E+3 | linear |
| 3 | 7.0E-2 | 4.1E-3 | linear |  | 13 | 1.3E-2 | 2.3E-2 | linear |
| 4 | 1.4E-2 | 2.0E+1 | linear |  | 14 | 2.2E-2 | 1.4E-3 | linear |
| 5 | 4.4E-2 | 1.0E-3 | linear |  | 15 | 2.7E-2 | 3.9E-3 | linear |
| 6 | 2.4E-2 | 4.8E+1 | linear |  | 16 | 2.9E-2 | 1.3E+1 | linear |
| 7 | 9.4E-3 | 4.8E+0 | linear |  | 17 | 7.1E-2 | 1.3E+1 | linear |
| 8 | 2.5E-2 | 1.5E-1 | linear |  | 18 | 1.1E-1 | 6.1E-2 | linear |
| 9 | 2.2E-2 | 6.6E-3 | linear |  | 19 | 3.5E-2 | 4.2E-2 | linear |
| 10 | 6.4E-3 | 1.0E+3 | linear |  | 20 | 2.9E-2 | 5.7E-1 | linear |
|  |  |  |  |  | mean | 4.0E-2 | 1.1E+2 |  |
|  |  |  |  |  | std | 2.9E-2 | 3.1E+2 |  |

**Table S14.** Comparison of minimalRNN with model filling strategy (RNN-MF) and SVM/SVR models with MFPCA. We observe that there was little performance difference between using MPFCA and linear interpolation (Table 4) for filling in missing data for the SVM/SVR models.

|  | mAUC<br>(more=better) | BCA<br>(more=better) | ADAS-Cog13<br>(less=better) | Ventricles<br>(less=better) |
| --- | --- | --- | --- | --- |
| RNN-MF | **0.944** ± 0.014 | **0.887** ± 0.024 | **4.30** ± 0.53 | **0.00156** ± 0.00022 |
| SVM/SVR (= 1tp)<br>(MFPCA) | 0.927 ± 0.013<br>(p = 0.011) | 0.837 ± 0.023<br>(p = $2.5 \times 10^{-7}$) | 5.19 ± 0.62<br>(p = $1.8 \times 10^{-4}$) | 0.00203 ± 0.00031<br>(p = $7.3 \times 10^{-5}$) |
| SVM/SVR (≤ 2tp)<br>(MFPCA) | 0.925 ± 0.013<br>(p = 0.002) | 0.834 ± 0.026<br>(p = $2.8 \times 10^{-6}$) | 5.22 ± 0.63<br>(p = $1.1 \times 10^{-4}$) | 0.00278 ± 0.00037<br>(p = $2.7 \times 10^{-7}$) |
| SVM/SVR (≤ 3tp)<br>(MFPCA) | 0.921 ± 0.013<br>(p = 0.001) | 0.824 ± 0.025<br>(p = $2.6 \times 10^{-7}$) | 5.46 ± 0.55<br>(p = $4.5 \times 10^{-7}$) | 0.00377 ± 0.00037<br>(p = $5.9 \times 10^{-7}$) |
| SVM/SVR (≤ 4tp)<br>(MFPCA) | 0.918 ± 0.012<br>(p = $2.2 \times 10^{-5}$) | 0.826 ± 0.019<br>(p = $4.1 \times 10^{-7}$) | 5.62 ± 0.58<br>(p = $9.4 \times 10^{-7}$) | 0.00436 ± 0.00035<br>(p = $1.2 \times 10^{-9}$) |
| SVM/SVR (= 1tp)<br>(Linear) | 0.929 ± 0.013<br>(p = 0.011) | 0.841 ± 0.023<br>(p = $2.5 \times 10^{-7}$) | 5.14 ± 0.62<br>(p = $1.8 \times 10^{-4}$) | 0.00199 ± 0.00031<br>(p = $7.3 \times 10^{-5}$) |
| SVM/SVR (≤ 2tp)<br>(Linear) | 0.926 ± 0.013<br>(p = 0.002) | 0.836 ± 0.026<br>(p = $2.8 \times 10^{-6}$) | 5.23 ± 0.63<br>(p = $1.1 \times 10^{-4}$) | 0.00230 ± 0.00037<br>(p = $2.7 \times 10^{-7}$) |
| SVM/SVR (≤ 3tp)<br>(Linear) | 0.923 ± 0.013<br>(p = 0.001) | 0.830 ± 0.025<br>(p = $2.6 \times 10^{-7}$) | 5.53 ± 0.55<br>(p = $4.5 \times 10^{-7}$) | 0.00261 ± 0.00037<br>(p = $5.9 \times 10^{-7}$) |
| SVM/SVR (≤ 4tp)<br>(Linear) | 0.919 ± 0.012<br>(p = $2.2 \times 10^{-5}$) | 0.832 ± 0.019<br>(p = $4.1 \times 10^{-7}$) | 5.68 ± 0.58<br>(p = $9.4 \times 10^{-7}$) | 0.00269 ± 0.00035<br>(p = $1.2 \times 10^{-9}$) |

## Footnotes

1 Although the goal is to (in principle) predict an unlimited number of time points into the future, the evaluation can only be performed using the finite number of timepoints available in the dataset.

